# The Origin and Evolution of Orphan Genes: A Case Study in Tea Plant Family

**DOI:** 10.1101/2024.02.01.578514

**Authors:** Lin Cheng, Yanlin Hao, Qunwei Han, Zhen Qiao, Mengge Li, Daliang Liu, Hao Yin, Tao Li, Wen Long, Shanshan Luo, Ya Gao, Zhihan Zhang, Houlin Yu, Xinhao Sun, Yiyong Zhao

## Abstract

Orphan genes and transcription factor genes (TFs) are pervasive across genomes, play pivotal roles as regulators in a myriad of biological processes. Despite their ubiquity, the evolutionary trajectories and functional divergence of these genes remain largely unexplored. Theaceae family, encompassing the economically and culturally significant tea plant, presents a unique opportunity to study these dynamics. Here, we decoded a nearly complete, chromosome-scale reference genome of *Stewartia gemmata* spanning 2.95 Gb. This study is enhanced by integrating the genome of *S. gemmata*, an early-diverging species within Theaceae, crucial for phylogenomic analyses and understanding the functional dynamics of orphan genes in this family. Our analysis confirmed the absence of a recent specific whole-genome duplication (WGD) event, with tandem duplications emerging as the predominant mechanism for gene duplication at ancestral nodes within Theaceae. By conducting an extensive comparative genomics analysis across 13 Theaceae and comparing these with a wide array of eukaryotic and prokaryotic proteins, we identified 37,618 orphan genes and 25,884 TFs in Theaceae. Interestingly, some orphan genes appear to have ancient origins in tea plant ancestors, suggesting relatively early origins with frequent gains and losses, conversely, many others seem more specific and recent. Notably, the orphan genes are characterized by shorter lengths, fewer exons and functional domains than TFs, implying relatively simpler functional roles. These orphan genes demonstrate diverse cellular localization and functions as predicted by GO/KEGG analysis, and are implicated in environmental response and flavor formation in tea plants. This study not only sheds light on the distinct evolutionary histories and functional divergences between orphan genes and TFs in Theaceae, but also contributes to our understanding of the genetic complexity and adaptability of this economically and culturally valuable plant family.

**Short summary:** The nearly complete genome of an early-diverging species *Stewartia gemmata* and phylogenomic studies provide insights into new gene evolution in Theaceae.

## Introduction

The family Theaceae, classified within the order Ericales, exhibits a remarkable level of biodiversity among angiosperms. With approximately 370 accepted species, the Theaceae family includes many economically, and horticulturally important species ^1^, such as the tea plant (*Camellia sinensis*), *Camellia oleifera*, *Camellia japonica*, *Camellia sasanqua*. Recent studies incorporating both morphological and molecular data have delineated three principal tribes within the Theaceae family: Theeae, Gordonieae, and Stewartieae. The Theeae tribe includes genera like *Camellia*, *Laplacea*, *Apterosperma*, *Polyspora*, and *Pyrenari*, while the Gordonieae tribe comprises *Gordonia*, *Franklinia*, and *Schima*. Finally, the Stewartieae tribe, which is recognized as a distinct and early-diverging lineage within the Theaceae family, encompasses the genera *Stewartia* and *Hartia*. Advanced phylogenetic analyses have provided insights into the evolutionary history of these tribes, suggesting that the most recent common ancestor (MRCA) of the Stewartieae tribe likely originated approximately 20.78 million years ago. ^2, 3, 4^. *Stewartia gemmata* is predominantly found in China’s southern provinces, including Hunan, Jiangxi, Fujian, Guangdong, and Yunnan, flourishing in mixed forests at altitudes of 900 to 1500 meters ^5, 6, 7^. *S. gemmata* typically reaches a height between four and eight meters, characterized by its smooth and greyish-yellow bark ^8, 9^. It is frequently used for ornamental purposes in horticulture owing to its vibrant flowers with high decorative and aesthetic appeal (Figure 1A). The bark, roots, and fruits of this plant, used in traditional medicine, have significant medico-economic value ^10^.

**Figure 1.**
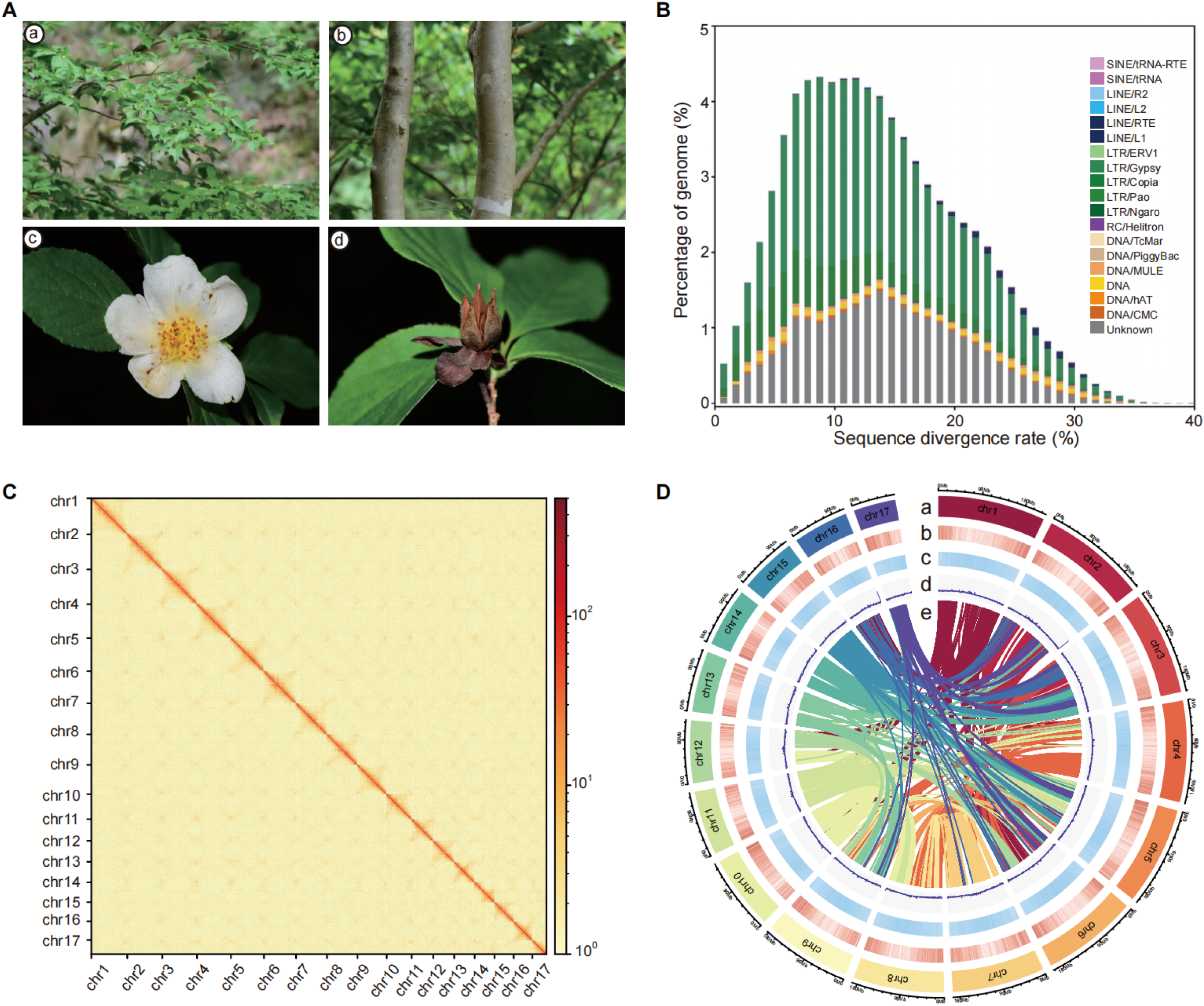
Morphological and genomic characteristics of *S. gemmata*. **A.** Morphological characteristics: Displaying various parts of *S. gemmata*. Panels a-d illustrate the leaf, stem, flower and fruit, respectively, showcasing the plant’s distinctive morphology. **B.** Transposon sequence divergence distribution: The graph depicts the divergence between transposon sequences in the *S. gemmata* genome and their corresponding sequences in the Repbase database. The x-axis represents the divergence rate, while the y-axis shows the percentage of transposon sequences in the genome at each divergence level. Different transposon types are distinguished by unique colors. **C.** Hi-C assisted genome assembly: Illustration of the chromosomal structure of S. gemmata as determined by Hi-C assisted genome assembly. The horizontal and vertical axes represent different chromosomes, highlighting the genome’s organization. **D.** Genomic annotation information: This section details various genomic features of *S. gemmata*, including a: Depiction of the 17 chromosomes. b: Visualization of gene density across the chromosomes. c: Distribution of repeated sequences. d: GC content variation. e: collinear connections between different chromosomes, indicating genomic synteny.

Orphan genes, also known as new genes or lineage-specific genes (LSGs), originate within the genomes of specific subsets of species ^11^. These genes have been extensively identified in various eukaryotes, such as plants ^12, 13, 14^, animals ^15, 16, 17, 18, 19, 20^, and fungi ^21, 22, 23^. As innovative genetic elements, they drive functional and phenotypic diversity and significantly influence the evolutionary processes of organisms^24, 25^. Studying LSGs relies on a stable and reliable phylogenetic foundation, with low copy nuclear genes being particularly effective in discerning phylogenetic relationships among angiosperms ^26, 27, 28^. Compared to morphological, anatomical, and cytological traits, as well as mitochondrial and chloroplast genomic markers, nuclear genes exhibit greater variability and longer sequence lengths. These characteristics enable nuclear genes to yield more detailed and comprehensive phylogenetic information, essential for understanding evolutionary histories and species diversification. ^29, 30^. Advances in genome sequencing have greatly facilitated the study of low-copy nuclear genes in plant phylogenetics ^31, 32, 33^. Analyses of genomes from basal species, especially those in early-diverging lineages, are crucial for clarifying phylogenetic relationships and accurately identifying LSGs. The genomes of basal angiosperms like *Amborella trichopoda* and *Nymphaea colorata* are instrumental in deepening our understanding of evolutionary mechanisms in early flowering plants ^34, 35^. In bamboo, genomic analysis has uncovered 1,622 LSGs, shedding light on their role in the plant’s rapid growth and offering insights into the genetic dynamics of bamboo stalk expansion ^36^. However, research on LSGs in the Theaceae family is limited by the scarcity of genomic data from basal species, significantly restricting the investigation into the LSGs’ contribution to tea flavor formation.

Whole genome duplication (WGD), a key evolutionary mechanism, has been documented to have taken place in the common ancestor of extant seed plants, angiosperms, and core Eudicots ^37, 38, 39^. WGD events have been identified in the early evolution of several major plant families, including Asteraceae, Brassicaceae, Fabaceae, Poaceae, and Rosaceae ^40, 41, 42, 43, 44^. Studies utilizing the genome of *Actinidia chinensis*, commonly known as kiwifruit, have suggested the occurrence of a WGD event, termed WGD-β, which was believed to be common to both *Actinidia* and *Camellia* species in Ericales ^45, 46^. Recent advancements in genomic analysis have illuminated the genomic localization of the WGD-β, predominantly associated with core Ericales, Primuloids, Polemonioids, and Lecythidaceae ^47^. Within the Theaceae family, genomic research primarily focuses on the genus *Camellia*, especially on *Camellia sinensis* (commonly known as tea plants) and oil-tea species ^48, 49, 50, 51, 52^. Recent investigations into WGD analysis in the Theaceae family have revealed discrepancies, particularly concerning the number of WGD events post whole-genome triplication-γ (WGT-γ) and the possibility of an independent WGD event within the family^39, 46, 49, 53, 54^.

Tea, a globally favored beverage, is produced from the leaves of *Camellia sinensis* (L.) O. Kuntze, a member of the Theaceae family ^53^. Its consumption is linked to various health benefits, such as preventing low-density lipoprotein oxidation, reduction of serum cholesterol, and decreased risk of cardiovascular syndromes ^55^. These benefits are largely due to tea’s rich composition of bioactive compounds, including tea polyphenols, theanine, and caffeine, which contribute to both its flavor and potential health advantages ^56^. The complex nature of tea flavor is influenced by environmental conditions, cultivation practices, processing methods, and the selection of specific tea cultivars ^57, 58, 59^. Whole genome duplication (WGD) events, characterized by the complete duplication of an organism’s genome, result in the introduction of additional copies of genes. This process significantly expands the pool of genetic material. Empirical evidence from prior research has demonstrated that WGD events contribute to increased genetic diversity within the tea plant (*Camellia sinensis*)^46, 49^. This increase in genetic diversity could potentially enhance the variation in alleles and genotypic compositions in tea plant ^60^. While research on the direct connection between WGD and tea flavor formation is limited, it’s plausible that WGD influences tea’s flavor profile through various mechanisms. Moreover, WGD can impact gene expression in metabolic pathways ^61^, including those responsible for the synthesis of flavor-related compounds. Changes in gene dosage resulting from WGD can lead to alterations in the production or accumulation of specific flavor compounds in tea leaves. In addition, polyploidy, which is the condition of having multiple sets of chromosomes, can confer certain advantages to plants, including increased vigor, adaptability, and stress tolerance ^62, 63^. While the WGD may contribute to the genetic foundation of tea flavor by generating novel genes through neofunctionalization of redundant duplicated genes, a comprehensive understanding of these relationships and underlying mechanisms requires further investigation ^64, 65, 66, 67, 68, 69^.

In our study, we successfully conducted a near-complete chromosomal-scale genome assembly of *S. gemmata*, a species that diverged early in Theaceae family by using whole-genome sequencing of short and long reads sequencing technologies. Our high-fidelity genome assembly revealed that tandem duplication is a pivotal mechanism in the evolutionary trajectory of Theaceae lineages and this is particularly notable given the absence of family-specific whole-genome duplications (WGDs) within Theaceae. We discovered 37,618 LSGs and numerous gene families within Theaceae that have undergone significant gene gain and loss events. Our comprehensive analysis, which involved comparing LSGs with more conserved TFs and performing GO/KEGG analysis, revealed the varied cellular localization of LSGs and their significant role in environmental response and flavor development in tea plants. These findings greatly enhance our understanding of gene dynamics within Theaceae, elucidating their impact on both biodiversity and the diversification of tea flavor.

## RESULTS

### Chromosome-level genome sequencing and assembly by integrating long-reads sequencing and Hi-C technologies

In preparation for the whole-genome sequencing of *S. gemmata*, flow cytometry was utilized to estimate its genome size, with maize and soybean genome serving as reference standards. This comparative analysis estimated the *S. gemmata* genome size around 2,722 Mb (Supplementary Fig. 1 and Supplementary Table 1). Subsequent genome survey analysis, using a Kmer value of 19, evaluated key genomic parameters including size, heterozygosity, and repeat sequence content. This analysis determined an estimated genome size of 2,739.02 Mb, a heterozygosity rate of 1.23%, and a repeat sequence content of 66.81% (Supplementary Fig. 2 and Supplementary Table 2), highlighting the genome’s notable complexity, particularly its significant heterozygosity exceeding 0.80%.

For sequencing the *S. gemmata* genome, a comprehensive approach was employed, integrating next-generation sequencing (NGS) by BGI (https://www.bgi.com/global), Oxford Nanopore long-read sequencing, and HiC-based chromatin mapping. The initial NGS phase yielded 303.88 Gb of raw data, refined to 302.53 Gb of clean data post-removal of joint and low-quality reads, with Q20 and Q30 values surpassing 90% (Supplementary Table 3). The Oxford Nanopore sequencing generated 155.03 Gb of raw data, culminating in 154.31 Gb of clean data after filtering (Supplementary Table 4). From these processes, 753 contigs with a combined length of 2,951.35 Mb were produced, with the longest measuring 41.33 Mb. The contig N50 value of 10.74 Mb affirmed the assembly’s enhanced continuity (Supplementary Table 5). Aligning NGS reads against the reference genome showed high mapping rate and coverage of 99.73% and 97.58%, respectively (Supplementary Table 6). BUSCO assessment with the embryophyta_odb10 dataset successfully assembled 1,573 of 1,614 genes, denoting 97.40% completeness (Supplementary Table 7). Furthermore, sequencing via the NovaSeq 6000 platform yielded 619.20 Gb of raw data, leading to 304 scaffolds spanning 2.95 Gb with an N50 value of 190.65 Mb (Supplementary Table 8). Hi-C assisted assembly facilitated the construction of 17 chromosomes, covering roughly 96.53% of the Hi-C assembled genome length of 2,848,999,739 bp (Supplementary Table 9). Notably, analysis indicated a stable transposon landscape recent evolutionary history, marked by an increased number of long terminal repeats (LTRs) during a transposon outbreak event in the genome of *S. gemmata*. A notable pattern was observed in the analysis, characterized by a single peak value at a sequence divergence rate of around 10 (Figure 1B). Post Hi-C assembly, the genome was accurately assigned to 17 chromosomes with a 96.53% anchoring rate, indicating uniform chromosome length distribution (Figure 1C, 1D). Gene density, repeat sequence density, GC content, and synteny analyses, depicted in Figure 1D, revealed higher gene density at chromosome ends compared to the middle regions.

### Annotation of genes and repeats for *S. gemmata* genome

An extensive annotation analysis was conducted on the genome of the *S. gemmata* to identify and characterize repeated sequences. The analysis indicated that a significant portion of the genome, approximately 2.37 Gb, consists of repeated sequences, accounting for about 80.18% of the total genome length (Supplementary Table 10). Notably, long terminal repeats (LTRs) represent 59.93% of the genome. Additionally, DNA transposons and RNA transposons were found to occupy 72.45 Mb (approximately 2.45% of the genome) and 1,806.50 Mb (around 61.21% of the genome), respectively (Supplementary Table 10). Using the GlimmerHMM dataset^70^, up to 253,374 genes were annotated. Moreover, a sequence homology-based prediction approach based on five closely related species annotated an average of 131,970 genes. This combined approach resulted in identifying a comprehensive set of 69,599 protein-coding genes (Supplementary Table 11). The average coding sequence (CDS) length was calculated to be 1,080.55 base pairs (bp), with genes comprising an average of 4.51 exons and an average exon length of 305.50 bp. The average intron length was found to be 2,289.50 bp. In comparison to related species such as *Camellia sinensis* var. *sinensis* (CSS) *‘*Tieguanyin’, *Camellia lanceoleosa*, *Camellia* DASZ, *Vaccinium darrowii*, and *Actinidia chinensi*s, the genes of *S. gemmata* exhibited a higher average number of exons (4.51 vs. 2.59) and exon length (305.5 bp vs. 260.71 bp). However, the average gene length of *S. gemmata* (9,418 bp) was significantly shorter than that in related species (21,286 bp) (Supplementary Table 11).

Kyoto Encyclopedia of Genes and Genomes (KEGG) pathway analysis revealed that 27.44% of the genes in *S. gemmata* genome showed homology with known molecular metabolic pathways. Additionally, Gene Ontology (GO) functional annotation identified 45,467 genes, representing about 65.33% of the total gene set. An extensive annotation analysis across multiple public databases indicated that a significant majority, 92.99%, of the genes had functional annotations (Supplementary Table 12). In the realm of non-coding RNAs, a total of 4,647 ncRNA genes were successfully annotated within the genome, this included the identification of 276 microRNAs (miRNAs). Moreover, the comprehensive analysis led to the annotation of 1,023 transfer RNAs (tRNAs), 679 ribosomal RNAs (rRNAs), and 2,669 small nuclear RNAs (snRNAs) (Supplementary Table 13).

### The resolution of Theaceae phylogeny through genome-scale phylogenomics analysis

Constructing a comprehensive phylogenetic framework is vital for understanding evolutionary dynamics, genetic variation intricacies, and genome architecture biology, especially in the context of whole genome duplication events across diverse genomes. To establish an extensive phylogenetic tree of Theaceae, we utilized datasets comprising 13 genomic sequences and 150 transcriptomic sequences from Theaceae, along with four genomic sequences as outgroups (Supplementary Table 14, 15). The 13 samples in our study, representing high-quality Theaceae genomes, included various *Camellia sinensis* varietals and other Theaceae species such as *Camellia oleifera* var. *Nanyongensis* and *Stewartia gemmata* (*Camellia sinensis* var. *sinensis* ‘Shuchazao’, *Camellia sinensis* var. *sinensis* ‘Longjing’, *Camellia sinensis* var. *sinensis ‘*Tieguanyin’, *Camellia sinensis* var. *sinensis* ‘Huangdan’, *Camellia sinensis* ‘DuyunMaojian’, *Camellia sinensis* var. *sinensis* ‘Biyun’, *Camellia sinensis* var. ‘YingHong9’, *Camellia sinensis* var. *assamica* ‘Yunkang 10’, *Camellia* DASZ, *Camellia lanceoleosa*, *Camellia oleifera* var. *Nanyongensis*, *Camellia chekiangoleosa*, and *Stewartia gemmate*). These were thoroughly analyzed to evaluate genomic data completeness, facilitating in-depth phylogenomic analyses and LSG detection to elucidate evolutionary relationships and genomic diversity within the family. Notably, all genome/transcriptome assemblies displayed BUSCO (C) scores over 50% (Figure 2) underscoring the high quality of the data used for the comprehensive phylogenetic analysis. Our comprehensive phylogenetic analysis offers robust insights into evolutionary relationships at both tribal and genera levels. The tribe Stewartieae emerged as sister groups to tribes Gordonieae and Theeae with maximal bootstrap support. The highly resolved phylogenetic reconstruction within Gordonieae and Stewartieae aligns remarkably with existing phylogenetic studies. Intriguingly, within the evolutionarily younger tribe Theeae, the oil-tea plants and the drinking-tea plants are discerned as sister groups. The wild tea plant (*Camellia* DASZ) emerged as the earliest diverging lineage among the drinking-tea plant species in Theeae. Furthermore, within the *C. sinensis* var*. assamica* (CSA) group, CSA ‘Yunkang10’ and CSA ‘Yinghong9’ were identified as successive sister groups to the CSS group (Figure 2).

**Figure 2.**
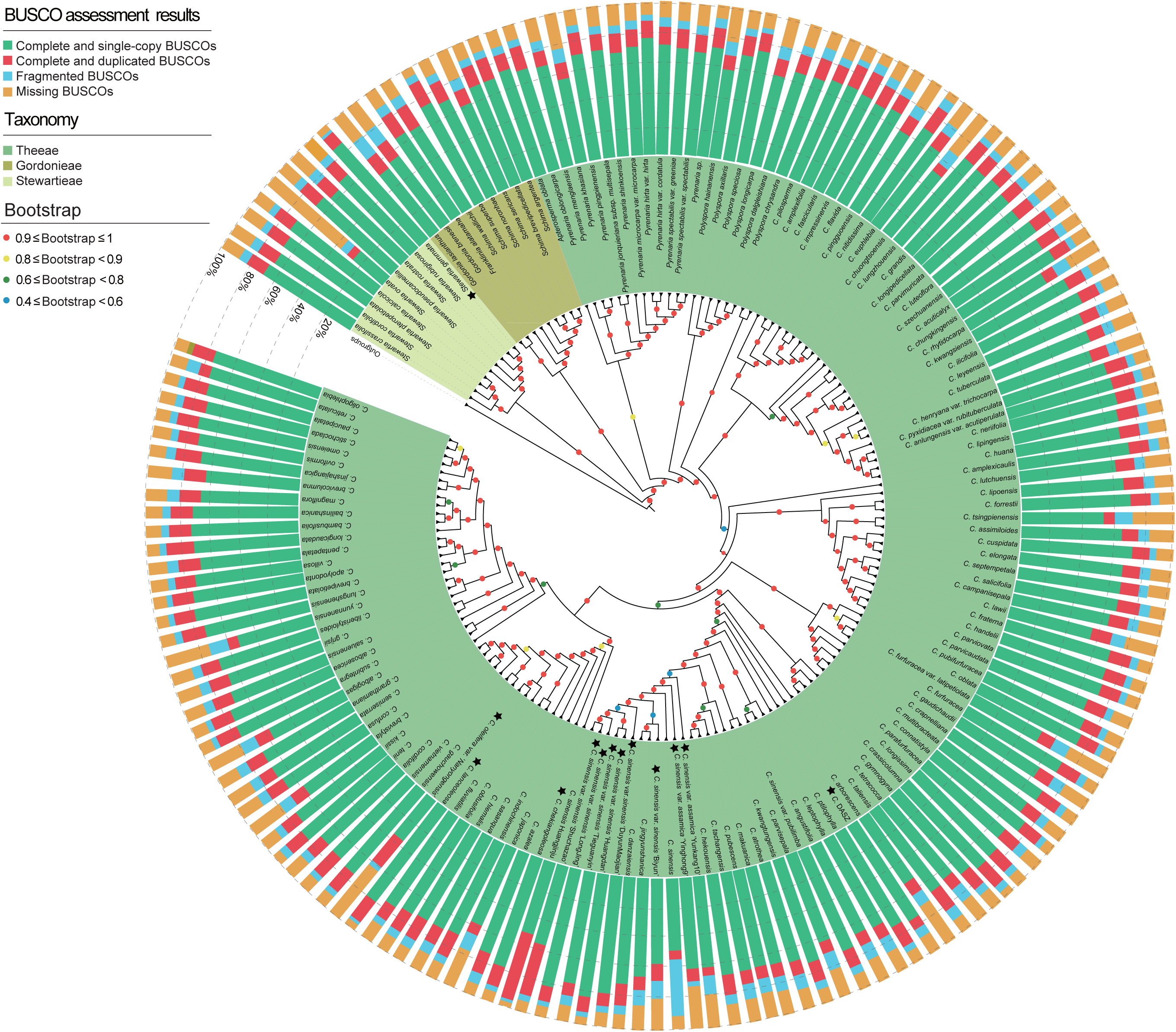
Phylogenetic relationship of Theaceae. The outer ring represents the BUSCO assembly results of the dataset, using the eudicots_odb10 reference database. The intermediate species names, marked by the letter C, signify the genus *Camellia*. Additionally, an asterisk highlights the 13 genome-level datasets specific to the Theaceae family. The inner circle comprises a series of differently colored dots, each indicating varying levels of support. A red dot denotes a support rating of 0.9 or higher, a green dot indicates a support rating between 0.6 and 0.8, and a blue dot signifies a support rating between 0.4 and 0.6.

For further comparative genomics analyses, we integrated our findings with the established Asterids phylogenetic framework, which features extensive species coverage at the order level^71^. Within this framework, Cornales and Ericales are identified as sister groups. Specifically, within Cornales, the Hydrangeaceae and Nyssaceae families are recognized as sister groups, highlighting their shared ancestry and close evolutionary relationship. In Ericales, Theaceae is positioned as the sister group to the most recent common ancestors of Roridulaceae, Actinidiaceae, Clethraceae, and Ericaceae families (Supplementary Fig. 3). Based on these results, we constructed a comprehensive species tree encompassing 31 genomes, which is detailed in Supplementary Fig. 3 with clearly labeled distinct nodes (Supplementary Table 15). This robust phylogenetic tree lays a solid foundation for investigating WGD events, identifying LSGs, elucidating evolutionary patterns, and understanding molecular changes within the Theaceae family.

### Evidence of an WGD in the Theaceae since WGT-γ

Our previous research highlighted a notable climate-related diversity rate shift in *Camellia*, which took place during the climate optimum period of the Miocene (MMCO)^3^. In this study, tricolpate pollen fossils were used as fossil calibration points for eudicotyledonous plant stem nodes. The divergence time of Theaceae was estimated to be around 69.40 Mya, while *Camellia* diverged approximately 38.77 Mya (Supplementary Fig. 4). A previous study conducted an analysis of the global climate trend over the past 65 million years (Figure 3A) ^72^. In our examination of the rate-through-time plot for Theaceae, we noted a significant increase in the speciation rate starting from the Core Tr. Theeae during the late Oligocene, around 30.8 million years ago (Mya) (Figure 3B) ^3^.

**Figure 3.**
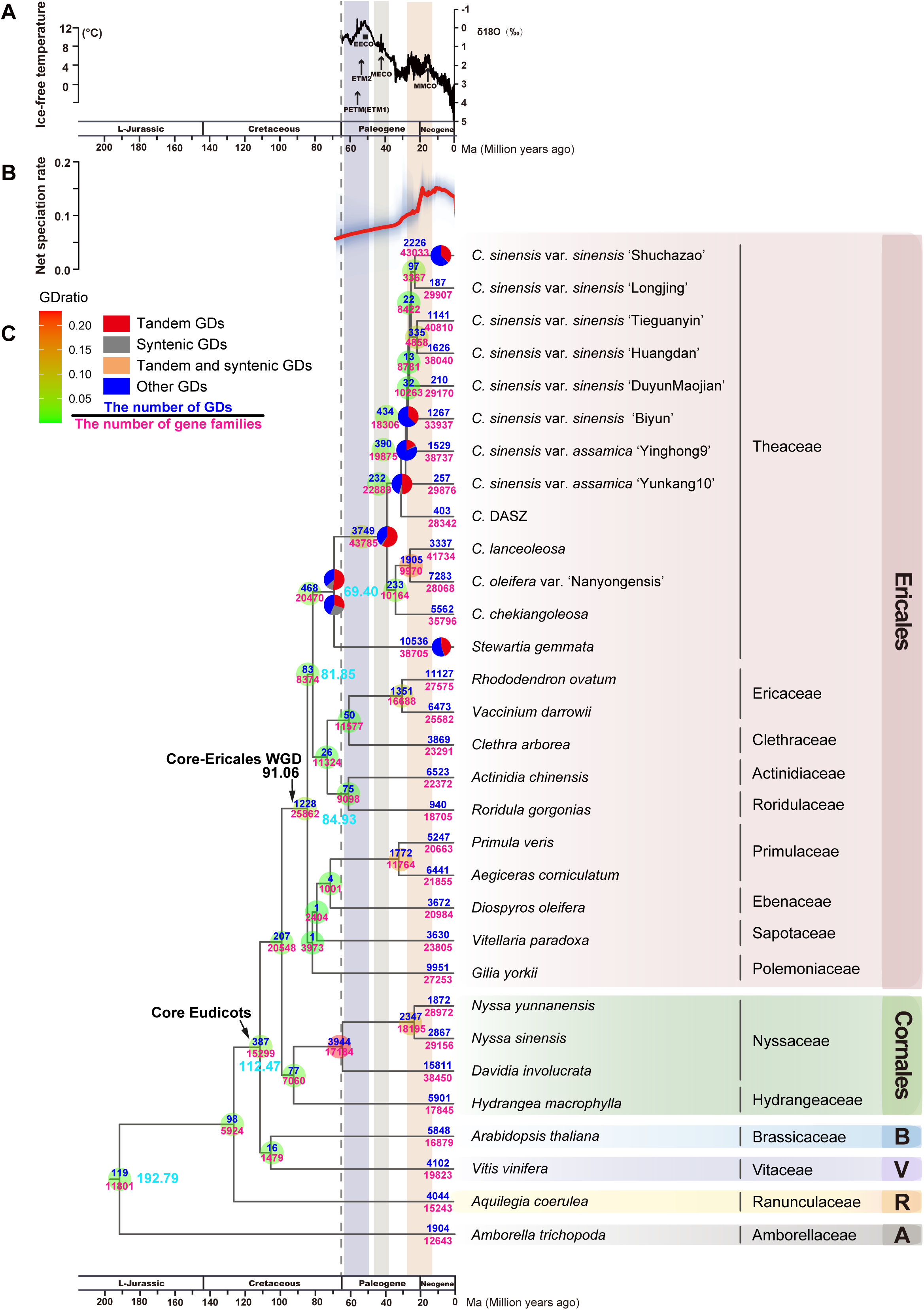
Divergence time of Ericales and identification of gene duplication events **in Theaceae.** **A.** Global temperature curve and climatic events: This panel illustrates the global temperature curve and significant climatic events over the last 65 million years. **B.** Diversification rate in Ericales: the rate-through-time plot of Ericales is dispayed here. The red line indicates the median diversification rate (species/milloon years), and the surrounding gray shadow represents the 95% credibility interval, offering insights into the rate of species diversification over time. **C.** Detection of gene duplication: This section focuses on the detection of gene duplication events in Theaceae, based on the gene tree-species tree. The cyan numbers adjacent to the nodes represent the differentiation times as predicted by the MCMCTree software. These provide an estimate of when each lineage diverged within the phylogenetic tree. Additionally, pie charts at the nodes illustrate the types of gene duplications observed. Each color in the pie chart represents a different duplication type: The top panel shows the red curve of global temperature change in Earth’s history based on data compiled from previous studies ^27^ and net speciation rate were adopted from our previous study^3^. Red: Proportion of gene duplications (GDs) involving tandem duplications; Gray: Proportion of GDs resulting from synteny, indicating duplications that arise due to genomic rearrangements preserving gene order; Orange: Proportion of GDs where both tandem and synteny duplications occur simultaneously; Blue: Proportion of GDs attributed to other types of duplications not classified as tandem or synteny. To the right of the species names, different families and orders are listed for quick reference. These include B for Brassicales, V for Vitales, R for Ranunculales, and A for Amborellales, aiding in the contextual understanding of the phylogenetic relationships and gene duplication events within these groups. B, Brassicales; V, Vitales; R, Ranunculales; A, Amborellales.

To investigate the occurrence of whole-genome duplication (WGD) events within the Theaceae family, we employed a comprehensive approach, utilizing gene and species tree reconciliation, genomic synteny assessment, and analysis of synonymous substitution rates (Ks). Our study revealed numerous GDs occurred at ancestral nodes of Theaceae. Specifically, at the node representing the most recent common ancestor (MRCA) of core Eudicots (N-27), we discovered 387 GDs. This finding echoes the well-documented whole-genome triplication-γ (WGT-γ) and subsequent WGD events within core Ericales (WGD-β), where we identified 1,228 gene duplicates at the Ericales node (N-21), as illustrated in Figure 3C and elaborated in Supplementary Figure 3. Further investigations within the Theaceae family unveiled a widespread occurrence of gene duplications across various nodes. At the MRCA of Theaceae (N-11), our analysis detected 468 gene duplications, representing 2.29% of the total gene families at that node. Remarkably, out of these duplications, only 130 gene duplicated pairs showcased a fully ABAB gene duplication pattern, comprising 27.78% of the gene pairs retained. A substantial number of duplications, 3,749 in total, were observed at the MRCA of *Camellia* (N-10), constituting 8.56% of the gene families present at that node. A predominant 80% of these duplications followed the ABAB pattern. This ABAB duplication trend was also present at the MRCA nodes of both cultivated tea (N-6) and the oil-tea plant (N-9), with the gene duplication (GD) ratios reflecting similar evolutionary patterns to those observed in other lineages within the genus *Camellia*.

In this study, intraspecific synteny analyses were conducted on *Camellia sinensis var. sinensis* ’Shuchazao’ and *S. gemmata*. The genome of *S. gemmata* was found to comprise 2,088 syntenic blocks distributed over 17 chromosomes. Dot plot analyses revealed prominent green syntenic blocks indicative of a 1:2 genomic correspondence in *S. gemmata* (Figure 4A). This observation corroborates the previously reported whole genome triplication event (WGT-γ) within the core Eudicots. Additionally, orange syntenic blocks demonstrated a 1:1 correspondence, aligning with recent whole genome duplication (WGD) events identified in the order Ericales (WGD-β) (Figure 4A). A similar synteny pattern was discerned in CSS ’Shuchazao’, with 1,909 syntenic blocks mapped to its 15 chromosomes (Supplementary Fig. 5A). These intraspecific synteny analyses suggests the absence of recent WGD events within the Theaceae family.

**Figure 4.**
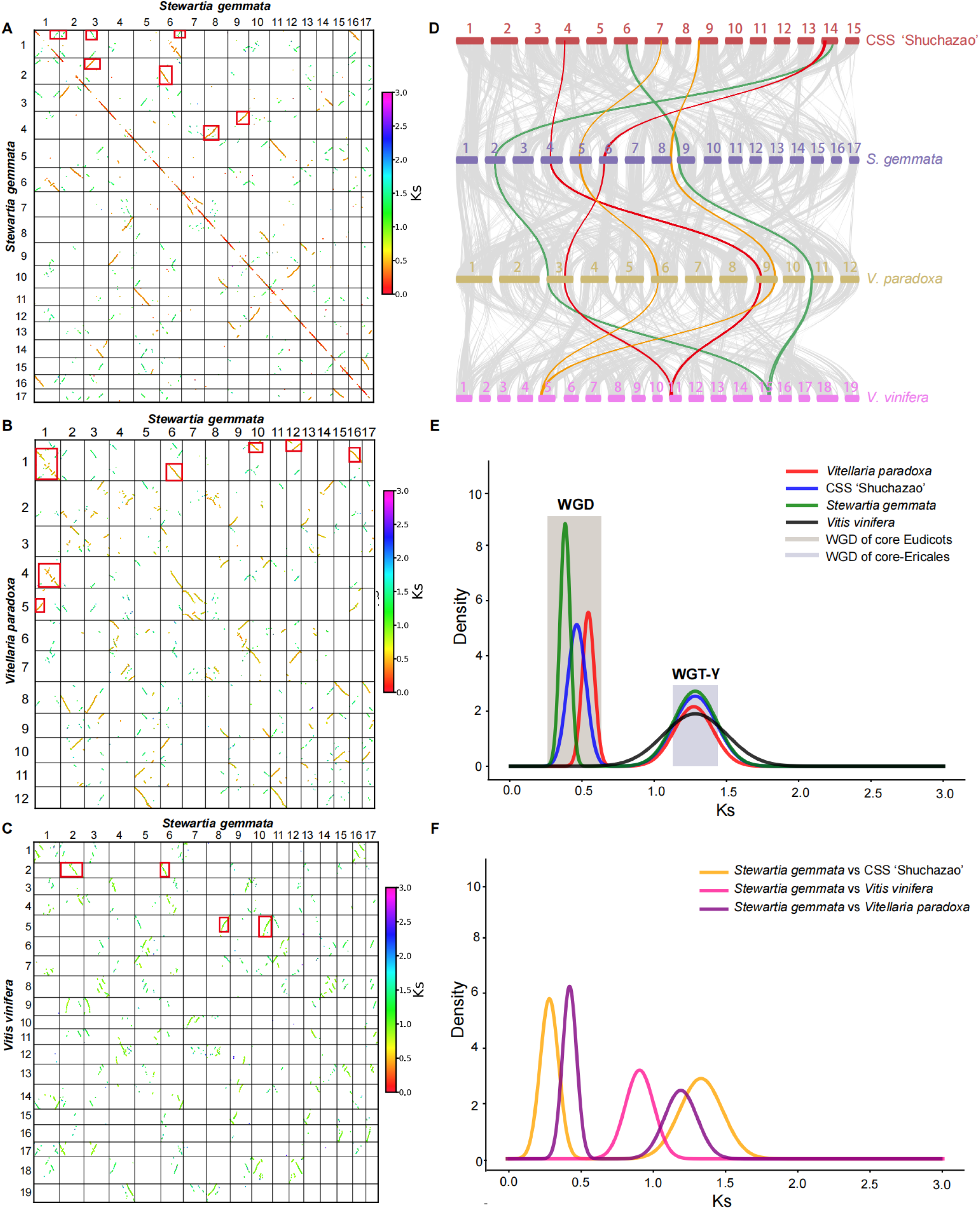
Identification of genome-wide duplication events in Theaceae. **A.** Dot-plot of syntenic blocks in *S. gemmata*: This panel displays a dot-plot representation of syntenic blocks within the *S. gemmata* genome, highlighting regions of genomic similarity and conservation. **B.** Synteny between *S. gemmata* and *V. paradoxa*: This dot-plot shows the syntenic relationships between the genomes of *S. gemmata* and *V. paradoxa*, illustrating the preserved gene order and genomic segments shared between these species. **C.** Synteny between *S. gemmata* and *V. vinifera*: similar to panel B, this dot-plot presents the syntenic blocks between *S. gemmata* and *V. vinifera*, indicating regions of conserved gene order. **D.** Conserved gene orders among four species: This illustration depicts the conservation of gene orders among *S. gemmata*, CSS *‘*Shuchazao’, *V. paradoxa*, and *V. vinifera*. It highlights a specific gene block in *V. vinifera* that is duplicated in the other three species, indicating a shared genomic heritage. **E.** *Ks* Distribution in paralogous genes: This graph shows the distribution of synonymous substitution rates (Ks) in paralogous gene pairs across genomes of *S. gemmata,* CSS *‘*Shuchaza*o’*, *V. paradoxa*, and *V. vinifera,* post-correction. This distribution helps in understanding the duplication events and their timing. **F.** Ks distribution in orthologous genes: similar to panel E, this graph displays the corrected Ks distribution in orthologous gene pairs among the four species. This analysis provides insights into the evolutionary relationships and divergence times between these species.

Furthermore, interspecific synteny analysis involving *S. gemmata* and other species, including CSS ’Shuchazao’, *Vitellaria paradoxa*, and *Vitis vinifera*, revealed syntenic block correspondences of 1:1, 1:1, and 2:1, respectively (Supplementary Fig. 5B and Figures 4B, 4C). The grape genome exhibited synteny with two chromosomes of *V. paradoxa*, denoting a 1:2 genomic correspondence. The synteny blocks of *S. gemmata*, CSS ’Shuchazao’, and *V. paradoxa* displayed a 1:1 correspondence on the corresponding chromosomes (Figure 4D), providing no evidence for WGD at the ancestral node of Theaceae. Post-divergence from *V. vinifera*, *S. gemmata* experienced an additional WGD event, which is shared with *Vitellaria paradoxa* and CSS ’Shuchazao’ and is congruent with the WGD detected in the core Ericales lineage.

Analysis of synonymous substitution rates (Ks) in paralogous gene pairs of *S. gemmata* (Figures 4E, 4F) revealed evidence of core-Ericales WGD and WGT-γ events, with discernible peaks at Ks values approximating 0.5 and 1-1.5, respectively (Figure 4E). The Ks values for paralogous gene pairs across *S. gemmata*, CSS ’Shuchazao’, and *V. paradoxa* ranged between 0 to 0.5 and 1 to 1.5. The distribution of these Ks values among paralogous gene pairs (Figure 4F) further supports the hypothesis of an absence of a recent, lineage-specific WGD event in the common ancestor of Theaceae.

### Identifications of gene duplication types in Theaceae

In the absence of whole genome duplication (WGD) events explicitly identified within Theaceae, an in-depth investigation was undertaken to elucidate the nature of gene duplication (GD) occurrences within this family. Our approach involved a meticulous gene tree-species tree reconciliation and an assessment of genomic synteny across various Theaceae taxa including *Camellia sinensis* var. *sinensis* ‘Shuchazao’ and *Stewartia gemmata*. The analysis revealed that a minority of GD pairs, which originated from ancestral Theaceae lineages at nodes N-11, N-10, N-7, and N-6, demonstrated synteny within their respective genomes. This observation suggests that the majority of these GD pairs were not generated through WGD events. Further scrutiny, quantifying the chromosomal proximity of these gene duplicates, indicated that gene pairs situated within a 10-gene radius comprised between 3.90% to 27.90% and 14.10% to 17% of the total duplicated gene pairs in CSS ‘Shuchazao’ and *S. gemmata*, respectively, as depicted in Supplementary Figure 6 and detailed in Supplementary Table 16. These findings underscore the significant role of tandem duplication events in the genesis of the gene duplication landscape observed in Theaceae.

### GO and KEGG enrichment analysis of tandem duplication genes

In order to explore the functional significance of tandem duplication in Theaceae, we conducted the Gene Ontology (GO) and Kyoto Encyclopedia of Genes and Genomes (KEGG) enrichment for those gene under tandem duplications. During the ancestral node of Theaceae, genes associated with CSS ‘Shuchazao’ underwent tandem duplication and enriched in active metabolic pathways such as zein and catecholoxidase (Supplementary Fig. 7). At the ancestral node of the *Camellia* genus, the genes with tandem duplication in CSS ‘Shuchazao’ were enriched in the defense responses, including oxidative stress response and biological stimuli, as well as flavonoid metabolism, terpene compounds, salicylic acid, and lignin synthesis (Supplementary Fig. 7). Tandem duplication genes, in addition to their fundamental role in sustaining life, play a crucial role in enhancing adaptability to the environment and contributing to species-specific traits. CSS ‘Shuchazao’ and *S. gemmata* exhibit distinct responses in tandem duplication genes to pollen recognition and defense mechanisms, with CSS ‘Shuchazao’ demonstrating responses to ultraviolet light and bacterial defense, while *S. gemmata* primarily shows fungal defense responses (Supplementary Fig. 7). Furthermore, CSS ‘Shuchazao’ responds to the metabolism of terpenes, fatty acids, and flavonoids at various nodes, enriching the spectrum of responsive enzymes. In contrast, *S. gemmata* is associated with the metabolism of terpenoids, vitamins, lignin, xylem and phloem development.

### Gene family expansion in Theaceae ancestors

In the evolutionary history of angiosperms, the gain and loss of gene families have been pivotal, shaping species-specific characteristics. Through PhyloMCL clustering of 1,245,359 protein sequences from 31 angiosperm genomes, we identified a total of 195,197 homologous gene clusters. These gene clusters were mapped to the species tree to analyze gene family expansion and contraction dynamics during the evolution of Theaceae (Supplementary Fig. 8). This analysis indicated that 2,350 gene families were gained at the MRCA of Theaceae. At the MRCA of the *Camellia* genus, a significant expansion to 7,306 gene families was observed (Supplementary Fig. 8). Additionally, 2,396 gene families were expanded at MRCA of cultivated teas, while 299 gene families were expanded at the MRCA of CSS (Supplementary Fig. 8). Moreover, at the MRCA of oil-teas including *Camellia oleifera*, *C. lanceoleosa* and *C. chekiangoleosa* with a total of 2,859 gene families were expanded. These expansions provide a pivotal insight into the genetic underpinnings of Theaceae, particularly in the context of environmental adaptation and the development of species-specific characteristics.

### Identification and distribution of LSGs and transcription factors (TFs) of Theaceae

In the Theaceae family, comprehensive genomic analysis led to the identification of 37,618 lineage-specific genes (LSGs), as detailed in Supplementary Table 17 and depicted in Supplementary Fig 9. A species-wise breakdown revealed diverse LSG counts: CSS ‘Shuchazao’ contained 3,071 LSGs, CSS ‘Longjing’ 966, CSS ‘Tieguanyin’ 1,671, CSS ‘Huangdan’ 4,253, CSS ‘Duyunmaojian’ 480, CSS ‘Biyun’ 1,067, CSS ‘Yinghong9’ 3,100, CSA ‘Yunkang10’ 1,668, C. DASZ 1,206, *C. lanceoleosa* 2,122, *C. oleifera* var. *Nanyong* 7,940, *C. chekiangoleosa* 4,610, and *S. gemmata* 5,464, as enumerated in Supplementary Table 18. LSGs constituted between 1% to 20% of the total gene count in these 13 Theaceae species, as illustrated in Figures 5A and 5B. Notably, over 40% of species-specific LSGs were discovered in *C. chekiangoleosa* and *S. gemmata*. In *C. oleifera* var. *Nanyong*, a significant number of LSGs, exceeding 3,016 gene families, were observed to be duplicated (Figure 5A). In contrast, the study identified a total of 25,884 transcription factor (TF) genes across the 13 Theaceae species, as documented in Supplementary Table 17. The distribution included 2,322 TFs in CSS ‘Shuchazao’, 1,875 in CSS ‘Longjing’, among others, with a comprehensive breakdown provided in Supplementary Table 19. The average TF gene count per species was approximately 2,000, showing a relatively stable presence, accounting for about 2%-6% of the total gene numbers in these plants, as shown in Figure 5B. This extensive analysis highlights the significant variation in the presence of LSGs across Theaceae species, contrasting with the more uniform distribution of TFs, thereby providing insights into the genomic complexity and diversity within this important plant family.

**Figure 5.**
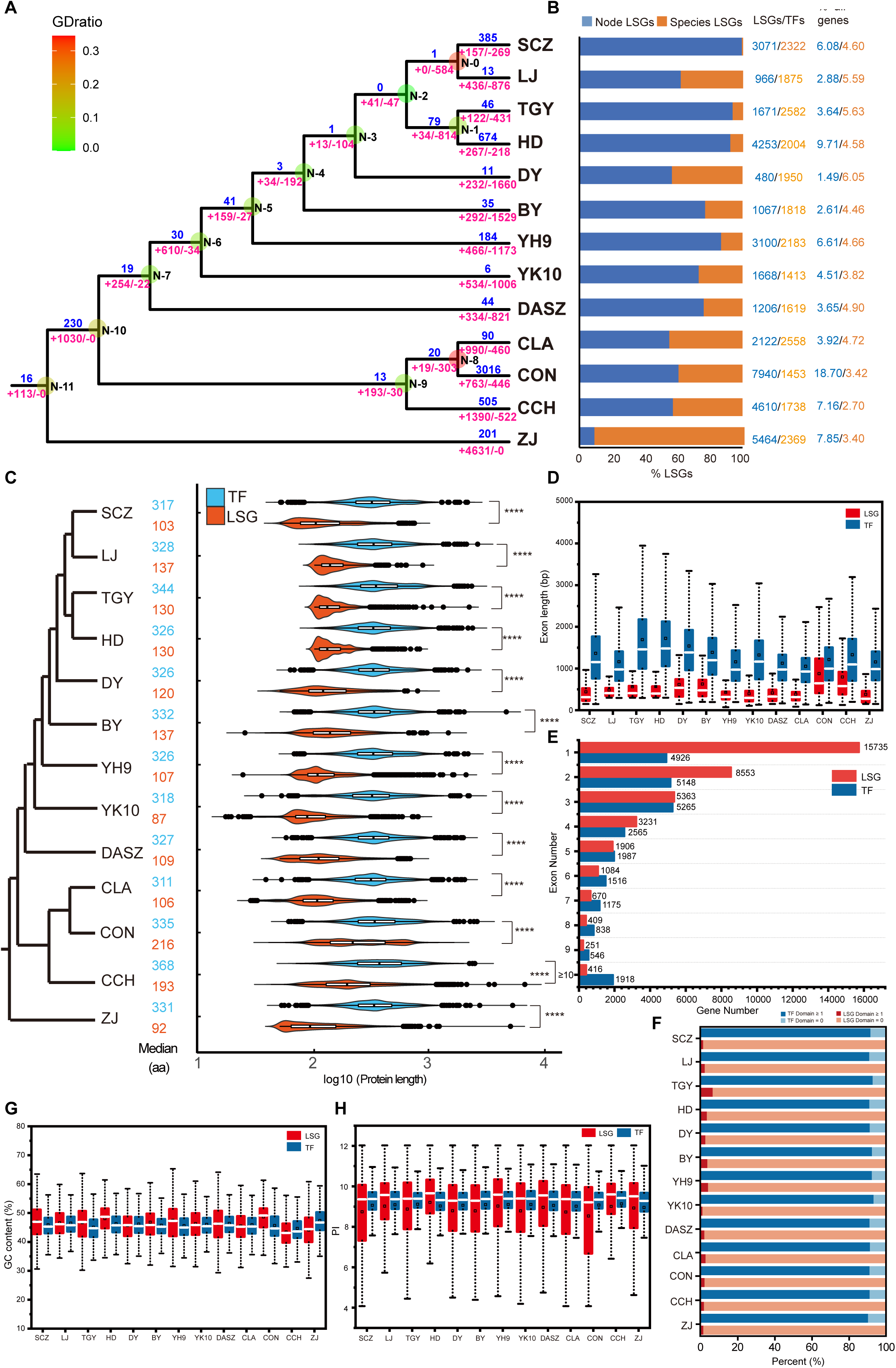
Distribution of lineage-specific genes and transcription factors in Theaceae. **A.** Gene duplication and family dynamics: This panel shows gene duplication (GD) counts above the horizontal line, with “+” symbols indicating gained gene families, “-” symbols for lost gene families, and “N” marking nodes in the evolutionary. **B.** Proportion of LSGs in ancestral nodes and species: The bar graph illustrates the proportion of lineage-specific genes (LSGs) in ancestral nodes and within individual species. The numbers on the right detail the count and percentage of LSGs and transcription factors (TFs) in each sample relative to the total gene count. **C.** Protein sequence length distribution: This section displays the distribution of protein sequence lengths for LSGs and TFs in 13 Theaceae plants. The left side presents phylogenetic relationships, the middle shows median protein sequence lengths for each group, and the right side depicts the distribution of protein length within each Theaceae plant. An independent sample t-test was performed, with \*\*\*\**P* < 0.0001 indicating statistical significance. **D.** Exon length distribution: This panel explores the exon length distribution of each lineage-specific gene and TF in 13 Theaceae plants. **E.** Number of exons per gene: The graph displays the distribution of the number of exons contained in individual LSGs and TFs, with the horizontal axis representing the number of genes and the vertical axis showing the number of exons per gene. **F.** Domain number distribution: This section shows the distribution of domain numbers in single LSGs and TFs across 13 Theaceae plants. The graph uses different colors to represent the proportion of genes with different domain counts, where dark blue indicates TFs with at least one domain, light blue for TFs without domains, dark red for lineage-specific genes with at least one domain, and light red for lineage-specific genes without domains. **G.** GC content distribution: This panel provides a comparison of GC content distribution between lineage-specific genes and TFs in 13 Theaceae plants. **H.** Isoelectric point (PI) distribution: The distribution of the isoelectric point (PI) for LSGs and TFs across the 13 Theaceae species is illustrated, offering insights into the biochemical properties of these proteins. Species Abbreviations: SCZ: *Camellia sinensis* var. *sinensis* ’Shuchazao’, LJ: *Camellia sinensis* var. *sinensis* ’Longjing’, TGY: *Camellia sinensis* var. *sinensis* ’Tieguanyin’, HD: *Camellia sinensis* var. *sinensis* ’Huangdan’, DY *Camellia sinensis* var. *sinensis* ’Duyun,’ BY: *Camellia sinensis* var. *sinensis* ’Biyun’, YH9: *Camellia sinensis* var. *assamica* ’Yinghong9’, YK10: *Camellia sinensis* var. *assamica* ’Yunkang10’, DASZ: *Camellia* DASZ, CLA: *Camellia lanceoleosa*, CON: *Camellia oleifera* var. *Nanyongensis*, CCH: *Camellia chekiangoleosa*, ZJ: *Stewartia gemmata*.

### Sequence characteristics of LSGs and TFs of Theaceae

In the 13 Theaceae plants, the LSGs exhibited shorter amino acid sequence lengths and exon lengths compared to the TFs, as shown in Figure 5C, D and detailed in Supplementary Table 20, 21. Contrasting with TFs, over 65% of LSGs in Theaceae had one or two exons. There were only 416 LSGs with more than 10 exons, while 1,918 TFs contained more than ten exons, as depicted in Figure 5E and Supplementary Table 20, 21. Additionally, a minimal proportion of LSGs encoded detectable functional protein domains, in stark contrast to over 90% of TFs, which possessed at least one protein domain (Figure 5F and Supplementary Table 22). In cultivated teas, LSGs generally showed higher GC content ratio than TFs (Figure 5G and Supplementary Table 20, 21). However, in *S. gemmata*, LSGs demonstrated lower GC content ratio compared to TFs (Figure 5G). The isoelectric points of most LSGs in Theaceae plants are higher than those of TFs, as indicated in Figure 5H and Supplementary Table 20 and 21. This comprehensive analysis elucidates distinct sequence characteristics between LSGs and TFs in Theaceae, underscoring the molecular diversity and complexity inherent in this plant family.

### Evolutionary characteristics of LSGs in Theaceae

In our study, we compared sequence characteristics at various ancestral nodes of Theaceae species, focusing on CSS *‘*Shuchazao’, CSA *‘*Yunkang10’, *C.* DASZ, *C. chekiangoleosa* and *S. gemmata*. Across the evolutionary timeline of these Theaceae plants, including *C.* DASZ, CSA *‘*Yunkang10’, and CSS *‘*Shuchazao’, there was a gradual reduction in sequence length, the amino acid length of LSG is getting shorter and shorter in the process of evolution. Consistent with this, LSGs in these representative species had fewer exons compared to TFs, aligning with the observations for amino acid sequence length. An analysis of GC content in LSGs and TFs at different evolutionary stages showed an initial decrease followed by a gradual increase in GC content, eventually approximating the levels in TFs across these species (Figure 6A). At the node N-11 of Theaceae, we identified 113 LSG families and 1149 TF gene families with TFs being about ten times more abundant. Across the 13 Theaceae plants, the number of LSG and TF gene families was generally around 1,000 (Figure 6B), indicating significant changes in the gain and loss of LSGs throughout evolutionary process. Our analysis revealed that at nodes N-10, N-9, N-7, N-6, and N-5, LSGs gained 1,030, 193, 254, 610, and 159 gene families, respectively, while TFs acquired fewer gene families at the same nodes. LSGs consistently gained at least five times more gene families compared to TFs. Conversely, at nodes N-8, N-4, N-3, N-1, and N-0, LSGs experienced a notable increase in lost gene families, surpassing the number lost by TFs by four times. Furthermore, LSGs within the 13 Theaceae plants exhibited a substantially higher count of both gain and lost gene families compared to TFs.

**Figure 6.**
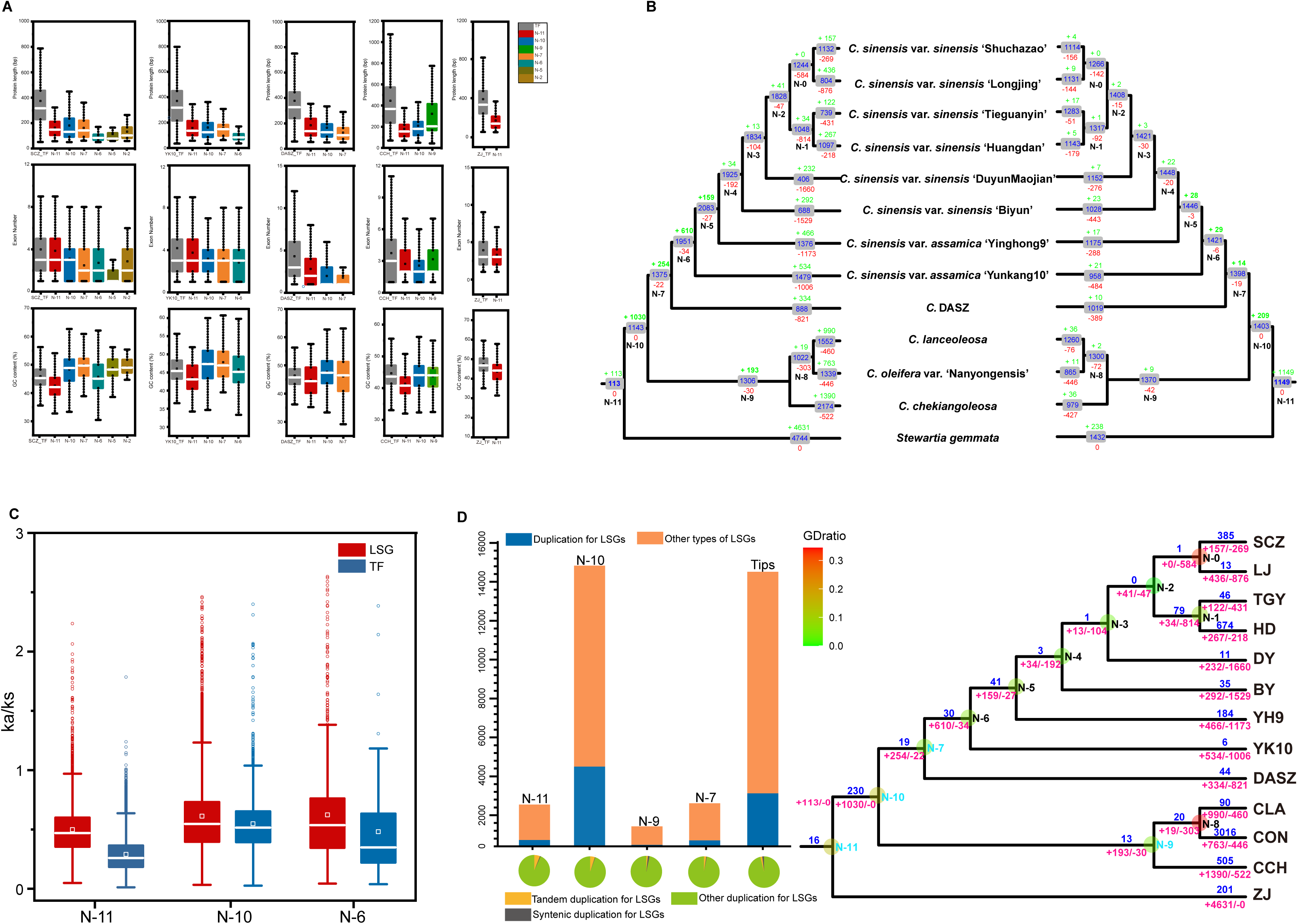
Evolutionary characteristics of lineage-specific genes in Theaceae. **A.** Protein sequence, exon number, and GC content distribution: This panel shows the distribution of protein sequence length, exon number, and GC content for lineage-specific genes (LSGs) and transcription factors (TFs) in CSS ‘Shuchazao’, CSA ‘Yunkang10’, *C.* DASZ, *C. chekiangoleosa*, and *S. gemmata* at different evolutionary nodes. Gray represents TFs identified in various Theaceae plants, while other colors indicate LSGs at different nodes. **B.** Gene family dynamics of LSGs and TFs: The figure illustrates the gene family gained and loss of LSGs and TFs in 13 Theaceae plants during their evolution. The left side represents gene family gains, while the right side shows losses. The numbers in the box indicate the count of gene families, with “+” for gains, “-” for losses, and “N” for nodes in the evolutionary tree. **C.** *Ka/Ks* values comparison: This section compares the ratio of nonsynonymous to synonymous substitutions (Ka/Ks) in orthologous gene pairs between LSGs and TFs at different nodes, namely N-11 (ancestor node of Theaceae), N-10 (ancestor node of the Camellia genus), and N-6 (ancestor node of cultivated tea). **D.** New gene duplication types: The right side shows the phylogenetic relationships of 13 Theaceae plants. Above the horizontal line, the number of gene duplications (GDs) in LSGs at ancestral nodes is indicated, with “+” denoting gained gene families and “-” representing lost gene families in LSGs. The bar chart on the left displays the number of LSG duplications and other types of LSGs at different nodes. Blue represents the number of LSG duplications, while orange indicates other types of LSGs. The pie chart demonstrates the proportion of LSGs in different duplication types: yellow for tandem duplication, gray for syntenic duplication, and green for other duplication types. “N” denotes nodes, and “Tips” indicate gene duplications occurring during the formation of the 13 Theaceae plants. Species abbreviations: The legend includes abbreviations for various Theaceae species studied: SCZ: *Camellia sinensis* var. *sinensis* ’Shuchazao’, LJ: *Camellia sinensis* var. *sinensis* ’Longjing’, TGY: *Camellia sinensis* var. *sinensis* ’Tieguanyin’, HD: *Camellia sinensis* var. *sinensis* ’Huangdan’, DY *Camellia sinensis* var. *sinensis* ’Duyun’, BY: *Camellia sinensis* var. *sinensis* ’Biyun’, YH9: *Camellia sinensis* var. *assamica* ’Yinghong9’, YK10: *Camellia sinensis* var. *assamica* ’Yunkang10’, DASZ: *Camellia* DASZ, CLA: *Camellia lanceoleosa*, CON: *Camellia oleifera* var. *Nanyongensis*, CCH: *Camellia chekiangoleosa*, ZJ: *Stewartia gemmata*.

We also compared the *Ka/Ks* ratios of orthologous gene pairs at various phylogenetic nodes, including the MRCA of Theaceae (N11), *Camellia* genus (N10), and the MRCA of cultivated tea (N6). *Ka/Ks* values from orthologous LSG pairs and TFs in *S. gemmata* were compared with eight representative species in cultivated tea groups, as well as CSS *‘*Shuchazao’, CSS *‘*Longjing’, CSS ‘DuyunMaojian’, CSS ‘Tieguanyin’, CSS ‘Huangtan’, CSS ‘Biyun’, CSA ‘Yinghong9’ and CSA ‘Yunkang10’, *C.* DASZ, *C. lanceoleosa, C. oleifera* var. *Nanyongensis,* and *C. chekiangoleosa* at N11 (Figure 6C, Supplementary Fig. 10A, D and Supplementary Table 23, 24). This method was similarly applied to the N10 and N6 (Figure 6C, Supplementary Fig. 10B, C, E, F). Results suggested that LSGs have undergone stronger positive selection, indicating higher evolutionary changes and potentially novel functional alterations. We observed a progressive increase in the *Ka/Ks* value distribution of homologous gene pairs of LSGs across different species between the nodes N-11 and N-6, suggesting an intensification of selection pressure on new genes throughout evolution.

Through GD analysis at various nodes, we identified 280, 4,078, 322, 124, and 2,758 LSGs have undergone GD events at their respective nodes, representing 13.41%, 28.37%, 14.52%, 11.60%, and 19.54% of the total number of LSGs (Figure 6D and Supplementary Table 25), which highlights the important role of GD in the production of LSG in Theaceae and its potential influence on species diversification. Further examination of the gene duplication types for LSGs showed that syntenic and tandem duplications contributed to less than 10% of the total number of LSGs generating from gene duplications (Figure 6D and Supplementary Table 25).

### Expression characteristics of LSGs and TFs in Theaceae

Gene expression data from CSS *‘*Shuchazao’, CSA *‘*Yunkang10’, *C.* DASZ, and *C. chekiangoleosa* were gathered across 21 different tissues and treatments (http://tpia.teaplants.cn/). For CSS *‘*Shuchazao’, the expression levels of LSGs tended to be lower than those of TFs (Figure 7A, 7B and Supplementary Table 26, 27). In CSA *‘*Yunkang10’, *C.* DASZ, and *C. chekiangoleosa*, 1,157, 938, and 3,043 LSGs were detected, representing 69.36%, 77.78%, and 66.01% of the total gene count, respectively (Figure 7C-E and Supplementary Table 28). For TFs, 1,400, 1,606, and 1,644 genes were expressed, constituting 99.08%, 99.20%, and 94.59% of the total TF count (Figure 7C-E and Supplementary Table 28). This demonstrates that over 60% of the LSGs in the four Theaceae species are expressed in at least one tissue or treatment (Supplementary Table 28).

**Figure 7.**
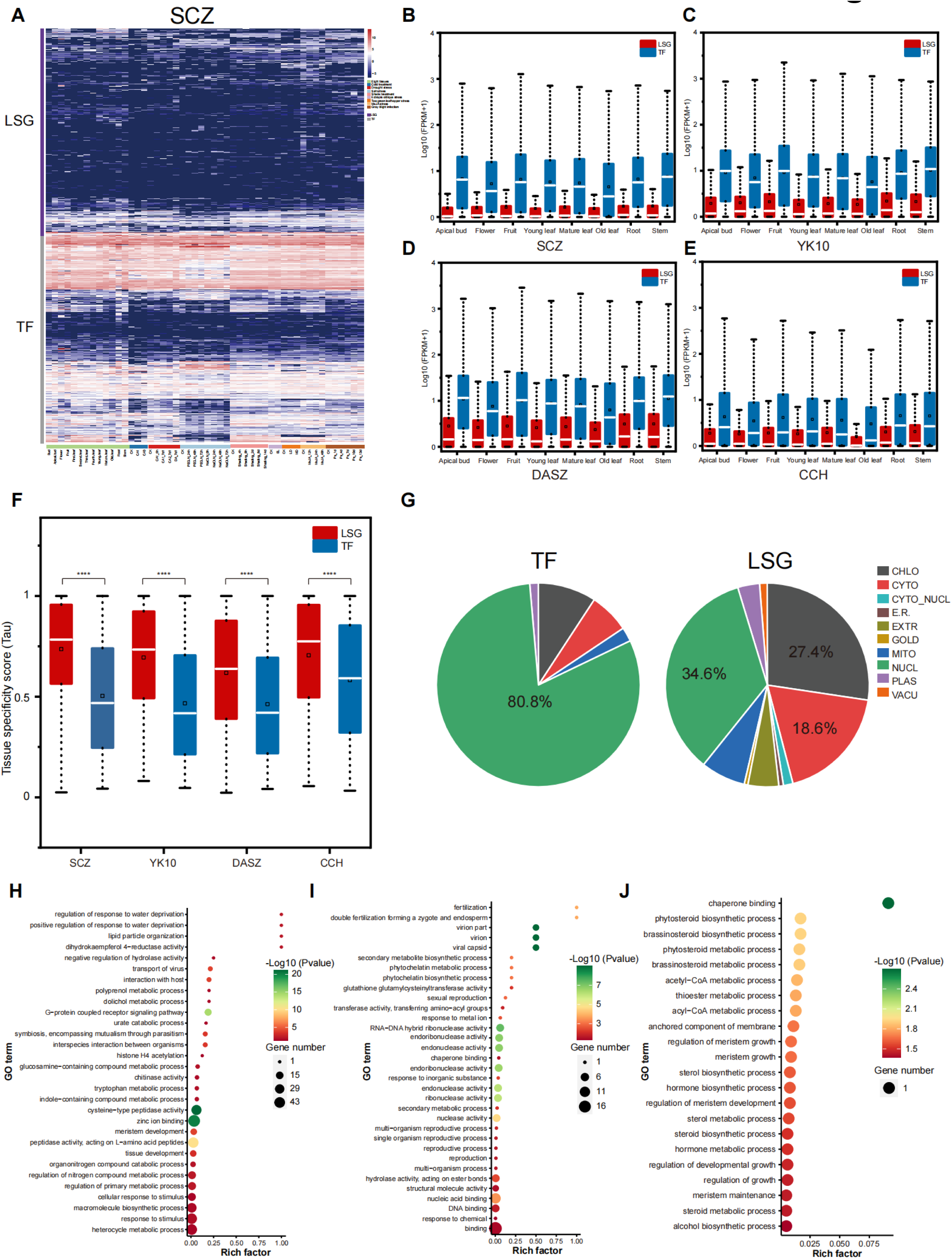
Expression characteristics and functional predictions of LSGs in Theaceae. **A.** Gene Expression in CSS ‘Shuchazao’: This panel shows the gene expression levels of lineage-specific genes and transcription factors (TFs) in CSS ‘Shuchazao’ across various tissues and under different environmental conditions. The horizontal axis lists different tissues and stress conditions including bud, flower, leaves of various ages, root, stem, and treatments like cold, drought, shading, pest stress, and infection. The vertical axis represents the expression levels of LSGs and TFs. **B-E.** Expression levels in various tissues: Panels B to E depict the expression levels of LSGs and TFs in eight different tissues for CSS ‘Shuchazao’ (B), CSA ‘Yunkang10’ (C), *C.* DASZ (D), and *C. chekiangoleosa* (E). These panels highlight the differential expression patterns of LSGs and TFs in key tissues. **F.** Tissue expression specificity index: This section presents the distribution of the tissue expression specificity index (Tau index) for CSS ‘Shuchazao’, CSA ‘Yunkang10’, *C*. DASZ, and *C. chekiangoleosa.* The Tau index quantifies gene expression specificity across tissues, with statistical significance indicated by an independent sample t-test (****P < 0.0001). **G.** Subcellular localization predictions: This panel predicts the subcellular localization of LSGs and TFs in Theaceae plants. The left and right sides represent TFs and LSGs, respectively, across 13 Theaceae plants. Various cellular components are indicated, including chloroplast (chlo), cytoplasm (cyto), cytoplasm_nucleus (cyto_nucl), extracellular (E.R.), golgibody (gold), mitochondria (mito), nucleus (nucl), plastids (plas), and vacuole (vacu). **H-J:** GO enrichment analysis: These panels present the Gene Ontology (GO) enrichment analysis of LSGs in *S. gemmata*, CSA ‘Yunkang10’, and CSS ‘Shuchazao’. The horizontal axis (rich factor) represents the ratio of differential genes under each metabolic pathway to all genes annotated in that pathway; the vertical axis indicates the enriched metabolic pathway. The size of each circle denotes the number of genes annotated to the corresponding GO item, and the color represents the significant level of enrichment results.

We analyzed the expression levels and breadth of LSGs across eight plant tissues in Theaceae, the findings showed that LSGs in CSS *‘*Shuchazao’, CSA *‘*Yunkang10’, *C.* DASZ, and *C. chekiangoleosa* generally had low expression levels (Figure 7A-E). To evaluate the expression breadth, we used a tissue specificity index value (τ), where higher τ values indicate stronger tissue-specific expression and lower values suggest broader expression. LSGs in these species typically had higher τ values compared to TFs (Figure 7F and Supplementary Table 29, 30).

Subcellular localization prediction for LSGs and TFs in Theaceae indicated that over 80% of the TFs were predicted to be nuclear, with a minority located in other compartments (Figure 7G and Supplementary Table 31). In contrast, less than 35% of LSGs were predicted to be nuclear, with the remainder anticipated to localize in various cellular areas, including the cytoplasm, chloroplasts, mitochondria, and extracellular matrix.

In *S. gemmata,* CSA *‘*Yunkang10’, and CSS *‘*Shuchazao’, LSGs contribute to growth, development and metabolic pathways (Figure 7H-J). In *S. gemmata*, some genes are involved in responding to external stimuli, regulating water shortage response, and nitrogen compound metabolism (Figure 7H). Notably, CSS *‘*Shuchazao’ differed by having genes annotated to biosynthetic pathways for phytosteroids, sterols, and brassinolides (Figure 7J). These findings suggest that LSGs may play a significant role in the adaptive capacity of Theaceae plants to diverse environmental challenges.

## Discussion

### High-quality genome of *S. gemmata* contributes to the resolution of phylogeny and provides a foundation for identifying LSGs within Theaceae

The acquisition of high-quality genomes from basal species is critically important for exploring the evolutionary dynamics and facilitating the identification of LSGs in tea plant family ^73, 74^. The rapid advancement of sequencing technology has facilitated the decoding of numerous plant genomes. Particularly notable is the sequencing of genomes from basal angiosperm species, including water lily and *Amborella trichopoda* ^34, 35^, has provided valuable insights into the early evolution of angiosperms and subsequent plant evolution. These genomic investigations have enhanced our understanding of the classification and functional roles of pivotal genes within angiosperms. Notably, the publication of the *Cercis chinensis* genome, a foundational species in the legume family, has provided a basis for investigating the origin and evolutionary models of new genes in legumes ^74^. To date, 12 genomes in Theaceae have been published, which primarily focus on two species: the tea plant and oil-tea plant ^48, 49, 50, 51, 53, 54, 75, 76, 77, 78^. The lack of genomic information on basal species in the Theaceae not only affects the resolution of the phylogenetic relationships within this family, but also hinders the identification of LSGs. The Stewartieae represents the earliest divergence clade within the Theaceae ^3^. In our study, we successfully assembled a high-quality genome for *S. gemmata*, a crucial reference genome for understanding gene family dynamics and the evolutionary origins of LSGs in ancestors of Theaceae and their sub-lineages. This accomplishment significantly enhances our understanding of genomic dynamics and evolutionary processes within this family. Our investigation focuses on analyzing the evolutionary divergence of LSGs compared to TFs and provides strong evidence for their functional roles in various developmental processes and their contribution to tea flavor formation.

### The absence of WGD events specific to Theaceae has been confirmed

Whole genome duplications (WGDs), prevalent in many land plant lineages known for high biodiversity, are believed to confer genetic novelties and complexities that enhance the adaptability of organisms to environmental challenges ^38, 40^. The Theaceae family, comprising three tribes and nine genera, has the *Camellia* genus encompassing about 50% of its species diversity ^79^. Although previous studies have commonly recognized that all Theaceae members underwent the core Eudicots whole-genome triplication (WGT-γ) event, the occurrence of subsequent WGD events in Theaceae, as well as the existence of their specific WGD events, remain topics of ongoing debate. Presently, there are three main viewpoints in this debate. One perspective suggests that following the WGT-γ event, Theaceae underwent a shared WGD-β event with kiwifruit and rhododendron ^46, 49, 54, 80^. The position of this WGD event is still highly controversial. The second viewpoint, emerging from a genomic study of CSS *‘*Shuchazao’, suggests an additional tea plant-specific WGD event that occurred ∼30 to 40 Mya ^53^, termed as Cm-α ^81^, alongside the WGT-γ and WGD-β events ^53, 81^. The third viewpoint posits that tea plants underwent a single tea plant-specific WGD event subsequent to the WGT-γ event, supporting the conclusion reached by Yang et al ^75, 82^. The challenge in resolving this debate is partly due to the limited availability of publicly accessible genomes within Theaceae, predominantly from the *Camelli*a genus. The utilization of tea plant as the sole representative species of Theaceae in most of the aforementioned studies led to inaccuracies in identifying WGDs at ancestral nodes of this family. Our study, incorporating the *S. gemmata* genome alongside CSS ‘Shuchazao’, *Vitis vinifera* and *Vitellaria paradoxa* genomes, aimed to investigate WGD events within the family. The *V. vinifera* genome has served as a valuable reference genome for evolutionary studies, primarily due to its ancestral Eudicot chromosome structure and the absence of additional WGD events, except for the WGT-γ ^83^. In previous studies investigating the WGD events in Theaceae, the genome of kiwifruit, the closest genus to *Camellia* in phylogenetic trees, was commonly used for *K*s analysis. However, compared to *V. paradoxa*, the kiwifruit experienced two subsequent tetraploidization events following the WGT-γ. These events include the *Actinidia* recent tetraploidization occurring approximately 18-20 Mya and the ancient tetraploidization dating back to around 50-57 Mya ^84, 85^. In addition, kiwifruit has undergone more substantial artificial selection as a significant fruit crop, which also resulting in a larger *Ks* peak value and faster evolutionary rate. This poses challenges for accurately identifying polyploidy events.

In our study, we selected *Vitellaria paradoxa* from the Sapotaceae family as the reference genome to explore evolutionary patterns, given its closer relationship to the ancestral node of core Ericales (Figure 3). We confirmed that Theaceae experienced only one round of WGD-β around 91.06 Mya since the WGT-γ, which is shared by other families in the order Ericales including Theaceae, Ericaceae, Clethraceae, Actinidiaceae, Roridulaceae, Primulaceae, Ebenaceae, Sapotaceae, Polemoniaceae, et al. (Figure 3, Figure 4, Supplementary Fig. 3). WGD has been widely acknowledged as a significant driving force in the evolution of speciation, adaptation, and diversification^86^. Emerging research suggests that polyploid plants with duplicated genomes exhibit enhanced adaptability and improved tolerance to diverse environmental conditions, which may have contributed to their higher survival rate during the Cretaceous-Tertiary extinction event about 66 Mya ^87, 88^. Our study indicates that despite the recent WGD event in Theaceae not aligning with the K-Pg boundary, the doubling of the genome played a crucial role in facilitating the differentiation and survival of Theaceae during the mass extinction event.

### New genes of Theaceae contribute greatly to the formation of tea flavor

Bioactive constituents like catechin, theanine, and caffeine are crucial in defining the unique properties and commercial value of tea plants ^89, 90^. Gene duplication events significantly influence the emergence of LSGs, with about 34.29% of LSGs in Theaceae attributed to gene duplications (Figure 6D and Supplementary Table 25). In our study, GO analysis showed that specific genes in *S. gemmata* are linked to response to external stimuli, positive regulation of water shortage responses, and participation in nitrogen compound metabolism. Nitrogen cycling is essential for synthesizing these key compounds in tea plants ^91, 92^. LSGs associated with fatty acid biosynthesis in the wild tea tree DASZ and within biosynthetic pathways of phytosteroids, sterols, and brassinolides in CSS ‘Shuchazao’ were identified (Figure 7J). Previous research has highlighted the correlation between fatty acid metabolism, tea leaf development, theanine synthesis, and tea quality ^93, 94, 95^. The developmental process of tea leaves is fundamental to understanding tea quality, suggesting these genetic elements may impact the complex biochemical pathways of flavor compound synthesis in tea plants.

These genes, involved in biosynthesis and modification of key flavor-related molecules, contribute to the diverse sensory characteristics of different tea varieties. Identifying and characterizing these genes provides insights into the diversification mechanisms of tea flavors. Furthermore, investigating the functional implications of LSGs opens avenues for targeted genetic manipulation and breeding strategies to enhance specific flavor traits in tea cultivars. Thus, exploring LSGs within the Theaceae family not only enriches our understanding of the molecular basis of tea flavor but also holds promise for the development of improved tea cultivars with tailored sensory profiles.

## Materials and Methods

### Plant material preparation and sequencing for *S. gemmata*

Sample collection and DNA extraction: fresh leaves were collected from a *S. gemmata* plant at Hangzhou Botanical Garden (Hangzhou City, Zhejiang Province; N30°16’, E120°12’, elevation 20-60 meters). Genomic DNA was extracted from these leaves using an enhanced CTAB method ^96^.

Next-generation sequencing library construction and sequencing: DNA samples underwent random fragmentation using the Covaris ultrasonic crusher. The Illumina sequencing library was prepared using the Nextera DNA Flex Library Prep Kit (Illumina, San Diego, CA, USA) following a series of steps including the DNA fragments with a target insertion fragment size of 150 bp were subjected to terminal repair, A-tail addition, sequencing splice addition, purification, and PCR amplification. After qualified library detection, high-throughput sequencing was performed on the DNBSEQ-T7 platform.

Third-generation sequencing library construction and quality assessment: large fragments of DNA (greater than 15Kb) were enriched and purified using magnetic beads. This process included the repair of damaged ends and polishing of the fragmented DNA. After purification, A-tailing was performed on both ends of the DNA fragments. Sequencing adapters from the SQK-LSK109 kit were then ligated to prepare the library. The constructed DNA library’s concentration was precisely measured using Qubit. Sequencing was conducted on the PromethION P48 sequencer (Oxford Nanopore Technologies, Oxford, UK) after loading the purified library onto R9.4 Spot-On Flow Cells.

Hi-C data sequencing: the sample was subjected to formaldehyde-induced cross-linking, followed by digestion with the restriction enzyme *Dpn*II (New England Biolabs, MA, USA). This step generated cohesive ends adjacent to the cross-linking sites. Subsequent steps included non-homologous end joining, circularization, DNA purification, capture, and library quality assessment. The final sequencing was performed on an Illumina Novoseq 6000 platform (Benagen Technologies, Hubei, China), generating paired-end reads of 150 base pairs each.

### Hi-C assisted chromosomal-level genome assembly and scaffolding

The raw data of Hi-C sequencing were processed using HICUP (v0.8.0) to extract valid interaction pairs ^97^. In the alignment process, reads that did not uniquely align to the reference genome were discarded. Additionally, invalid read pairs and duplicates resulting from PCR amplification were filtered out. This filtering step ensured only valid interaction pairs were retained for subsequent analysis. The initially assembled contigs were scaffolded using 3D-DNA and Juicer software ^98, 99^, leveraging these valid interaction pairs to refine the draft genome sequence. This methodology enabled the effective anchoring of contigs to chromosomes, culminating in the assembly of the *S. gemmata* genome at the chromosomal level.

### Genome annotation of *S. gemmata*

Employing a *de novo* approach, the software RepeatModeler was used to predict model sequences based on the genome sequence from the *S. gemmata* genome ^100^. Additionally, LTR_FINDER software was utilized to predict LTR (Long Terminal Repeat) sequences^101^. The LTR_retriever software was utilized to conduct a de-redundancy process on sequences predicted by LTR_FINDER ^102^. The RepeatMasker subroutine RepeatProteinMask was used for predicting transposable element (TE) protein type repeat sequences. Gene structure prediction was performed using a comprehensive approach, combining transcriptome prediction, homology prediction, and *de novo* prediction.

For the next and third generation sequencing data, the genome was compared using the software HISAT2 v2.1.0 and minimap2 v2.17 ^103, 104^. The aligned data in BAM format were further processed using Stringtie v2.1.4 with the parameter “-a p15” to reconstruct transcripts ^105^. TransDecoder v5.1.0 software (https://github.com/TransDecoder/TransDecoder) was utilized to predict coding frames within sequence. Homologous predictions involved five related species including CSS ‘Tieguanyin’, *C.* DASZ, *C. lanceoleosa*, *A. chinensis*, and *V. darrowi*i, with protein sequences compared against the genome using the tBLASTN algorithm. Results were further utilized for transcript and coding predictions with Exonerate v2.4.0 (https://github.com/nathanweeks/exonerate). Gene predictions were performed using Augustus v3.3, Genscan v1.0, and GlimmerHMM v3.0.4, with Genscan (http://genome.ucsc.edu/cgi-bin/hgTrackUi?g=genscan) specifically used for the prediction process ^106, 107, 108^. The MAKER v2.31.10 software integrated and consolidated gene sets from these methods.

Sequence and motif similarities were used to annotate the gene function of *S. gemmata*. Functional information and metabolic pathway associations of protein sequences were analyzed using diamond BLASTp v2.0.11.149 ^109^ against databases such as UniProt, NR, and KEGG ^110^. Protein domain architecture and transmembrane regions were predicted using InterProScan v5.52-86.0, querying protein sequences against secondary databases within InterPro, including CDD, Gene3D, Hamap, Panther, Pfam, Phobius, Pirsf, Pirsr, Prints, Prosite, Sfld, Smart, Superfamily, and Tig. Hmmscan v3.3.2 was used for domain prediction to identify conserved sequences, motifs and domains of proteins ^111^. tRNAscan-SE v1.23 (parameter -q) was utilized to search for tRNA sequences based on their structural characteristics ^112^. ncRNA sequences were annotated with INFERNAL v1.1.2 with the parameters “-cut_ga -rfam -nohommonly - cpu 15” using Rfam database ^113^.

### Integration of genomic and transcriptomic datasets: assembly, annotation and **BUSCO completeness assessment**

In this study, we compiled a comprehensive collection of 181 publicly available angiosperm datasets, comprising 31 genomic and 150 Theaceae transcriptomic datasets. This collection, augmented by our high-quality genome assembly of *S. gemmata,* is crucial for the accurate identification and characterization of new genes. The genomic data included 13 high-quality datasets from Theaceae and 18 genomes from outgroups taxa such as Ericaceae, Clethraceae, Actinidiaceae, Roridulaceae, Primulaceae, Ebenaceae, Sapotaceae, and Polemoniaceae. Additionally, we incorporated genomes from Nyssaceae and Hydrangeaceae (Cornales) ^114, 115, 116, 117^, Brassicaceae (Brassicales) ^118^, Vitaceae (Vitales) ^83^, and *Aquilegia coerulea* ^119^ as a representative of basal Dicotyledons. The genome of *A. trichopoda*, representing basal angiosperms, was also included ^34^. Transcriptomic data mainly comprised 133 samples from the tribe Theeae, 9 samples from the Tr. Gordonieae, and 8 samples from the Tr. Stewartieae. Transcriptomic data were from the National Center for Biotechnology Information (NCBI, https://www.ncbi.nlm.nih.gov).

The software Trimmomatic (v0.39) ^120^ was used to perform filtering on a dataset consisting of 149 publicly available transcriptomes and shallow genome data samples (referred to as “Raw data”) (Supplementary Table 14). *Camellia sinensis* var ‘Huangjinju’ is represented by a CDS and PEP file that was directly downloaded. The process involved removing low-quality reads using parameters as “LEADING:10TRAILING:10 SLIDINGWINDOW:4:20 MINLEN:36”. *De novo* assembly of all transcriptomes was conducted with Trinity (v2.11.0) ^121^, constructing contigs from the raw transcriptomic data (Grabherr et al., 2011). The software TransDecoder version 5.5.0 (http://transdecoder.sourceforge.net/, accessed in August 2023) was utilized for the prediction of coding sequence (CDS) regions. CD-HIT program (v4.8.1) was employed to reduce redundancy within each assembly, with a parameter of “-c 0.98” ^40, 122, 123, 124^. The software BUSCO (v5.2.2) was used to assess the completeness of gene annotations for each sample using the eudicots_odb10 database, specifically designed for eudicot plant species ^125^. Details regarding the transcriptomes generated in this study (Supplementary Table 14), as well as the assembly completeness assessment using BUSCO, are provided in Supplementary Table 14, 15.

### Phylogenetic analysis

We used our customized script to convert the nucleic acid sequences into its corresponding amino acid sequences. For a comprehensive protein comparison, the software diamond was utilized to perform all against all BLASTp analysis on the amino acid sequences ^109^, with a default E-value 1E-5 for significance. The Markov Clustering Algorithm (MCL, https://micans.org/mcl/) v14.137, was employed to cluster the gene pairs obtained from the BLASTP comparison ^126^. This algorithm facilitated the grouping the gene pairs into distinct clusters based on their similarity bitscores for each paired homolog.

Multiple sequence alignment of the clustered amino acid sequences was conducted using MAFFT v7.487 ^127^. The aligned amino acid sequences were then converted into nucleic acid sequences using PAL2NAL v13 software ^128^. To remove poorly aligned regions and enhance alignment quality, trimAl v1.2 software was employed with the parameters “automated1” to improve the overall alignment quality.

For phylogenetic analysis, IQ-TREE v2.1.4-beta software ^129^ was used to construct a maximum likelihood (ML) gene family tree. The best model for tree construction was automatically determined by the software ModelFinder ^130^ for accurate phylogenetic inference, and 1000 bootstrap replicates were performed to validate the robustness of tree topology. Finally, ASTRAL-Pro ^131^ was employed to reconstruct the coalescent species tree.

### Estimation of the divergence time and origination of whole genome duplication events in the MRCA of Core Ericales

For divergence time estimation, we selected two fossil calibration points and two secondary calibration points as divergence time markers. The fossil calibration points chosen were 125-247.2 million years (Myr), representing the crown group node of Angiosperms ^132^ and the earliest tricolpate pollen fossils, marking the stem node of Eudicots around 125 Myr ^132^. The secondary calibration points were 79.8-102.5 Myr for the stem node of Theaceae and 39.6-74.7 Myr for the crown node of Theaceae ^133^. The MCMCTree program within PAML software was utilized for molecular clock analysis, using 1,030 orthologous groups (OGs) with at least 90% coverage and gene lengths over 800 bp ^134^. This analysis incorporated the auto-correlation model, the General Time Reversible (GTR) site substitution model, and prior probabilities.

Markov Chain Monte Carlo (MCMC) sampling estimated the posterior distribution of node ages, discarding the initial 200,000 generations and sampling every 20 generations over a total of 500,000 generations. Convergence was checked by repeated analysis and examining a sufficient sample size.

The ages of WGDs detected in this study were estimated based on the assumption of a constant rate of synonymous mutation accumulation. If a WGD event is flanked by two species divergence events in the species tree, the upper limit of the WGD age (denoted as *T^prio^*^r^) was set at the time of species divergence preceding the WGD, and the lower limit (denoted as *T^pos^*^t^) at the divergence time following the WGD. The emergence times of lineages on the species tree, derived from a recent study using fossil data and computational estimates of chloroplast genes^135^, determined *T^prior^*and *T^post^* for each WGD event. The time of a specific WGD event, *T^wgd^*, was determined using the following function:

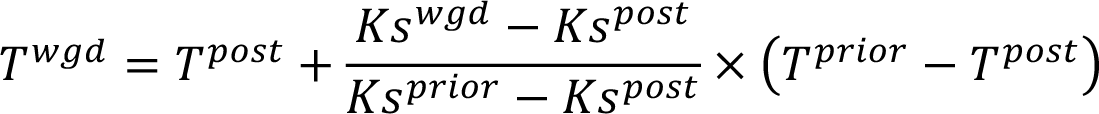

In the context of WGD events*, Ks* values play a critical role. *Ks^prior^* and *Ks^post^* refer to the highest synonymous substitution (Ks) values observed between orthologs of two diverging lineages before and after the WGD event, respectively. These values were derived by conducting all-against-all BLAST analysis to compare reciprocal best matched genes between paired species. Conversely, *Ks^wgd^*represents the average Ks value across all gene duplicates resulting from the specific WGD event being studied. **Detection of gene duplication events in different Theaceae lineages**

For predicting gene duplication events, we conducted gene and species tree reconciliation using the Tree2GD tool ^136^. This approach allowed us to reconciliate gene tree data with species tree, facilitating the identification of gene duplication events within the evolutionary context. Additionally, we assessed collinearity both between and within species using the WGDI software with default parameters ^137^. This software analyzes genomic collinearity, which is crucial for understanding the conservation and diversification of gene order among different species or within a same species over evolutionary time.

### Synonymous substitution rates (*Ks*) correction

We performed the *Ks* correction to accurately determine the origin of core-Ericales WGD by a previous described method^138^. This approach assumes normal distribution for the evolutionary rate *Ks* values, species A was selected as the reference to correct the *Ks* values of species B.

If 𝑋_*_ ∼ 𝑁(𝜇_*_, 𝜎_*_^+^) represents the distribution of *Ks* values in species A, and the duplicated gene pairs in species B follow 𝑋, ∼ 𝑁(𝜇,, 𝜎,^+^) , then the correction coefficient is given by:

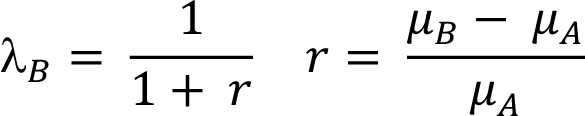

The corrected value of 𝑋,, denoted as:

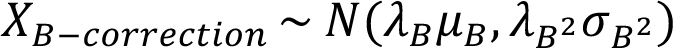

If there is another species C, there is:

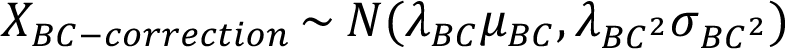

### Identification of new genes and transcription factors (TFs) in Theaceae

In this study, the CDS sequences of 31 data sets were translated into amino acid sequences by the guidance of codon code. These amino acid sequences were subjected to “All-Against-all” comparison using Diamond software with an E-value threshold of 1e^-5^. PhyloMCL was employed to cluster the genes by analyzing the based BLASTp bitscores, resulting in the identification of orthologous clusters or gene families across the 31 data sets. The dolloparsimony module within the Tree2GD software package was utilized to identify gene family gain and loss events. Gene families obtained from ancestral nodes, internal branches, and individual species were subsequently considered as candidate gene sets for specific genes in Theaceae. Candidate genes underwent sequential comparison against multiple databases including nonredundant protein sequences (NR), nonredundant nucleotide sequences (NT), SWISS-PROT (a manually annotated and reviewed protein sequence database), 1000 Plant transcriptomes initiative (OneKP) databases, and a collection of 150 Theaceae transcriptomic assembly gene sets. Candidates not matching any entries in these databases were designated as Theaceae LSG set one. This comparison process involved species classification, exclusion of non-Theaceae genes, and rejection of matches with Cyanobacteria. Additionally, the retrieved genes needed to meet sequence similarity criteria across all databases, with a threshold of at least 70% similarity, and each gene required a minimum of five matches with other genes (Supplementary Fig. 9). These criteria helped form gene set two. The cumulative total of these two gene sets represents the new genes identified within Theaceae.

### Analysis of gene sequence characteristics of Theaceae

The LSGs of 13 samples within Theaceae were analyzed, focusing on their proportion within the total gene count of each species. Various characteristics such as amino acid sequence length, exon number and length, and GC content were determined. The isoelectric point (PI) of the encoded proteins was calculated using the software TBtools^139^. Additionally, the Pfam database was employed to identify protein domains within the LSGs and TFs of Theaceae, and the number of protein domains was ascertained. Data visualization was performed using the software Origin for effective presentation of the findings ^140^.

To ascertain the reliability and precision in identifying lineage-specific genes (LSGs), we employed a comprehensive approach incorporating public high-quality Theaceae genomes, including basal species *Stewartia gemmata*, sequenced and assembled in this study. Additionally, we integrated genome data from 18 representative outgroup species, selected based on data quality, to enhance the robustness of our analysis. In the process of identifying LSGs in Theaceae, a total of 192,751 genes from 56,150 gene families were identified, originating from ancestral nodes, internal branches, and the species themselves. These genes formed the candidate gene pool for LSGs in Theaceae (Supplementary Fig. 9, and Figure 5A).

### Analysis of gene expression characteristics of Theaceae

To access expression data of CSS *‘*Shuchazao’, CSA *‘*Yunkang10’, *C.* DASZ, and *C. chekiangoleosa* at different developmental stages and habitats, we used the Tea Plant Information Archive (TPIA) online website (http://tpia.teaplants.cn/). For predicting the subcellular localization of TF protein sequences specific to Theaceae, the WoLF PSORT online website tool (https://wolfpsort.hgc.jp/) was employed. Additionally, to investigate the gain and loss of LSGs and TFs within the Theaceae gene family evolution, the Tree2gd software was utilized ^136^.

The tissue expression specificity index, known as the Tau index (τ), was employed to quantify the extent of gene expression across various tissues ^141^. The Tau index (τ) is calculated from normalized expression levels of genes (𝑥_)_) in different tissues relative to the maximum expression level observed across all tissues. The Tau index (τ) ranges between 0 and 1, with 0 indicating broad expression across tissues and 1 highly indicating tissue-specific expression. The formula for the calculation of Tau index (τ) is as follows:

Here, N represents the total number of tissues analyzed. This index provides a quantitative measure of gene expression specificity, aiding in the understanding of gene function and regulation in different tissue types.

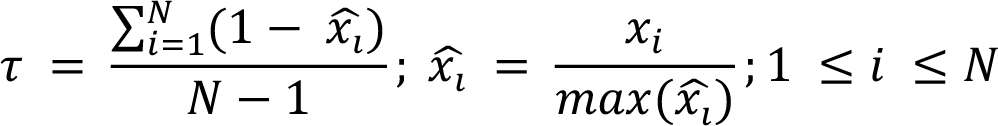

### Gene selection pressure analysis

To compare LSG and TF orthologous gene pairs at different nodes, multiple sequence alignments were conducted using the MAFFT software ^127^. This alignment provided the basis for subsequent analyses of the evolutionary dynamics between these species. For computing the comparison results of the matched orthologous gene pairs, the *KaKs*_Calculator tool was utilized ^142^. This tool applied the NG (Nei-Gojobori) method to determine the ratio of *Ka* to *Ks* substitutions^142^. This ratio, known as the *Ka*/*Ks* ratio, is a critical indicator of evolutionary pressure on genes; a *Ka/Ks* ratio greater than 1 suggests positive selection, less than 1 indicates purifying selection, and a ratio around 1 implies neutral evolution.

## Acknowledgments

We thank to Dr. Xinxin Zhu for providing the image materials of *S. gemmata*. We are particularly thankful for the anonymous reviewers’ valuable comments on the manuscript. We are grateful to the State Key Laboratory of Big Data, Guizhou University for providing super computing services for data analysis. This work was funded by the Program for Science & Technology Innovation Talents in Universities of Henan Province (HASTIT, No. 21HASTIT040) and the Scientific and Technological Project in Henan Province (212102110447). Additional support was provided by the Key Laboratory of Functional Agriculture in Higher Education of Guizhou Province, China, designated by the grant number Qian Jiao Ji [2023]007.

## Author details

^1^Henan International Joint Laboratory of Tea-oil tree Biology and High-Value Utilization, College of Life Sciences, Xinyang Normal University, Xinyang 464000, China. ^2^Guizhou Key Laboratory of Functional Agriculture, College of Agriculture, Guizhou University, Guiyang 550025, China. ^3^State Key Laboratory of Public Big Data, College of Computer Science and Technology, Guizhou University, Guiyang 550025, China. ^4^Xinyang Normal University Library, Xinyang Normal University, Xinyang 464000, China. ^5^Broad Institute of MIT and Harvard, Cambridge, MA 02142, USA ^6^College of Science, Northeastern University, Boston, MA 02115, USA

## Author contributions

Yiyong Zhao and Lin Cheng conceived and designed the research and experiments; Yiyong Zhao, Zhihan Zhang, Lin Cheng, Yanlin Hao, Mengge Li prepared samples and subsequent sequencing. Yanlin Hao and Lin Cheng performed most of the bioinformatic analyses under the assistance from Yiyong Zhao. Zhen Qiao, Qunwei Han, Mengge Li, Daliang Liu, Hao Yin, Tao Li, Ya Gao participated in analyzing the data. Wen Long provided support for the Linux cluster server. Shanshan Luo provided support for the data curation and storage, respectively. Critical revisions were made by Yiyong Zhao, Lin Cheng, Yanlin Hao, Houlin Yu and Xinhao Sun. All authors thoroughly reviewed and endorsed the final version of the manuscript.

## Conflict of interest

The authors declare no competing interests.

All data that support the findings of this study are available from the corresponding author upon a reasonable request.

**Supplementary Fig. 1. Genome size estimation of *S. gemmata* via flow cytometry**. R1 denotes the flow pattern peak of *S. gemmata*, R2 corresponds to the peak flow pattern of *Glycine max*, and R3 and R4 reresent the composite flow patterns of the *Glycine max* intranuclear replication peak and *S. gemmata*, respectively.

**Supplementary Fig. 2.** Kmer analysis in *S. gemmata* at Kmer = 19. This figure illustrates the distribution of Kmer depth and the frequency of Kmer species at Kmer = 19.

**Supplementary Fig. 3. Phylogenetic analysis of 31 Angiosperm plants.** The C in species name stands for *Camellia*, the N signifies the node, and the vertical line on the right indicates the family and order affiliations of each species.

**Supplementary Fig. 4. Estimation of species divergence times by MCMCtree.** The blue bars at the nodes indicate 95% confidence intervals. The geological timescale, presented at the bottom, is measured in millions of years, with “L-Jurassic” referring to the Late-Jurassic period. On the right of each species name, families and orders are denoted, with “B” for Brassicales, “V” for Vitales, “R” for Ranunculales, “A” for Amborellales.

**Supplementary Fig. 5.** Dot-plot of syntenic blocks wthin CSS ‘Shuchazao’ (A) and the dot-plot between CSS ‘Shuchazao’ and *S. gemmata* (B).

This figure represents a comparative syntenic block dot plot between the genomes of CSS ‘Shuchazao’ (A) and *S. gemmata* (B).

**Supplementary Fig. 6.** Analysis of duplicated gene pairs in CSS ‘Shuchazao’ (A) and *S. gemmata* (B). The bar chart displays the counts of duplicated genes in CSS ‘Shuchazao’ and *S. gemmata* across various nodes and within the species themselves. Blue bars indicate duplicated genes on the same chromosome, yellow for genes duplicated due to synteny, and red for other gene duplications. The pie chart shows the distribution of distance between duplicated gene pairs on the same chromosome, with 0-10 denoting a distance of up to ten gene intervals.

**Supplementary Fig. 7.** GO and KEGG enrichment analysis of tandem duplicated genes in *Camellia sinensis* var. *sinensis* ‘Shuchazao’ and *S. gemmata*. In the GO enrichment analysis, N-11 represents the ancestral node of Theaceae, and N-10 signifies the ancestral node of the *Camellia* genus. “SCZ” and “ZJ” are abbreviations for *Camellia sinensis* var. *sinensis* ‘Shuchazao’ and *Stewartia gemmata*, respectively. Circle size in the plots corresponds to the number of genes annotated to each GO term, while the color denotes the significance level of enrichment, with a Q-value < 0.05. In the KEGG enrichment analysis, the numbers on the bar graph indicate the count of genes enriched in each pathway.

**Supplementary Fig. 8.** Gene family dynamics in 31 Angiosperm species. The values in the boxes represent the number of gene families, with “+” indicating a gain and “-” a loss of gene families.

**Supplementary Fig. 9.** The pipeline for the identification of LSGs in Theaceae.

This figure delineates the methodology used in identifying lineage-specific genes (LSGs) within the Theaceae family.

**Supplementary Fig. 10.** Evolutionary rate analysis of LSGs and TFs in Theaceae, *Camellia* genera, and cultivated teas. Distribution of Ka/Ks values of LSG pairs (A) and transcription factors (D) at the ancestral node of Theaceae, *Stewartia gemmata*, with *Camellia chekiangoleosa*, *Camellia oleifera* var. ‘Nanyongensis’, *Camellia lanceoleosa*, *Camellia* DASZ, CSA ‘Yunkang10’, CSA ‘Yinghong9’, CSS ‘Biyun’, CSS ‘Huangdan’, CSS ‘Tieguanyin’, CSS ‘Duyun’, CSS ‘Longjing’ and CSS ‘Shuchazao’, respectively. Distribution of Ka/Ks values of LSG pairs (B) and transcription factors (E) at the ancestral node of *Camellia* genus, *Camellia chekiangoleosa*, with *Camellia* DASZ, CSA ‘Yunkang10’, CSA ‘Yinghong9’, CSS ‘Biyun’, CSS ‘Huangdan’, CSS ‘Tieguanyin’, CSS ‘Duyun’, CSS ‘Longjing’ and CSS ‘Shuchazao’, respectively. Distribution of Ka/Ks values of LSG pairs (C) and TFs (F) at the ancestral node of cultivated tea, CSA ‘Yunkang10’, with CSA ‘Yinghong9’, CSS ‘Biyun’, CSS ‘Huangdan’, CSS ‘Tieguanyin’, CSS ‘Duyun’, CSS ‘Longjing’ and CSS ‘Shuchazao’, respectively. “CSS” is *Camellia sinensis* var. *sinensis* and “CSA” is *Camellia sinensis* var. *assamica*.

**Supplementary Table 1.** Genome size estimation of *Stewartia gemmate* using flow cytometry.

Note: “Count” refers to the number of nuclei analysis. “Mean” represents the average fluorescence value, indicative of DNA content in the nucleus. “CV” denotes the coefficient of variation. R1 signifies the peak pattern of *Stewartia gemmate*, R2 for G*lycine max*, and R3 and R4 are the combined flow patterns of the G*lycine max* intranuclear replication peak and *Stewartia gemmate*.

**Supplementary Table 2.** K-mer analysis results from next-generation sequencing of *Stewartia gemmata* genome.

Note: “X” indicates the depth of sequencing in the analysis.

**Supplementary Table 3.** Quality metrics from next-generation sequencing of *Stewartia gemmata* genome.

Note: “Q20” and “Q30” represent the percentage of total bases with Q_score values exceeding 20 and 30, respectively.

**Supplementary Table 4.** Data from third-generation sequencing of the *Stewartia gemmata* genome.

**Supplementary Table 5.** Genome continuity assessment of *Stewartia gemmata* genome.

Note: “N50” and “N90” describe specific contig lengths at which the sum of lengths of the longest contigs comprises 50% and 90% of the total length, respectively. “Average” is the mean length of contigs. “Median” is the median contig length of contigs. “Min” is the minimum length of contigs. “Max” is the maximum length of contigs.

**Supplementary Table 6.** Consistency assessment of *Stewartia gemmata* genome. Note: “map_rate” denotes the comparison rate of next-generation sequencing data.

**Supplementary Table 7.** Integrity assessment of *Stewartia gemmata* genome.

**Supplementary Table 8.** Genome assembly data statistics for *Stewartia gemmata*. Note: “N50” and “N90” describe specific scaffold lengths at which the sum of lengths of the longest scaffolds comprises 50% and 90% of the total length, respectively. “Average” is the mean length of scaffolds. “Median” is the median scaffold length of scaffolds. “Min” is the minimum length of scaffolds. “Max” is the maximum length of the scaffolds.

**Supplementary Table 9.** Chromosome length statistics of *Stewartia gemmata* genome.

**Supplementary Table 10.** Analysis of repetitive sequences in *Stewartia gemmata* genome.

**Supplementary Table 11.** Gene structure prediction information of *Stewartia gemmata.*

**Supplementary Table 12.** Functional annotation of *Stewartia gemmata* genome-encoded genes.

**Supplementary Table 13.** Non-coding RNA annotation statistics in *Stewartia gemmata* genome.

**Supplementary Table 14.** Assembly and integrity assessment of transcriptome data from 150 Theaceae plants.

**Supplementary Table 15.** Completeness assessment of genomic data from 31 Angiosperm plants.

**Supplementary Table 16.** Analysis of gene duplication types in *Camellia sinensis* var. *sinensis* ’Shuchazao’ and *Stewartia gemmata*.

**Supplementary Table 17.** Identified LSGs and TFs in 13 Theaceae plants.

**Supplementary Table 18.** Distribution of LSGs in 13 Theaceae plants.

Note: Refer to Supplementary Table 15 for the mapping of three-letter abbreviations to species names.

**Supplementary Table 19.** Distribution of TFs in 13 Theaceae plants.

Note: Refer to Supplementary Table 15 for the mapping of three-letter abbreviations to species names.

**Supplementary Table 20.** Protein characteristics of LSGs in 13 Theaceae plants.

**Supplementary Table 21.** Protein characteristics of TFs in 13 Theaceae plants.

**Supplementary Table 22.** Protein domain profile of LSG-encoded proteins and TFs in 13 Theaceae plants.

**Supplementary Table 23.** Ka/Ks values of orthologous LSG gene pairs in Theaceae.

**Supplementary Table 24.** Ka/Ks values of orthologous TF gene pairs in Theaceae.

**Supplementary Table 25.** Gene duplication types of LSGs in 13 Theaceae plants.

**Supplementary Table 26.** Expression profiles of LSGs in *Camellia sinensis* var. *sinensis* ’Shuchazao’.

Note: Expression data are sourced from Tea Plant Information Archive (TPIA, http://tpia.teaplants.cn/).

**Supplementary Table 27.** Expression profiles of TFs in *Camellia sinensis* var. *sinensis* ’Shuchazao’.

Note: Expression data are derived from Tea Plant Information Archive (TPIA, http://tpia.teaplants.cn/).

**Supplementary Table 28.** The number of genes with detectable expression in LSGs and TFs within *Camellia sinensis* var. *sinensis* ’Shuchazao’, *Camellia sinensis* var. *assamica* ’Yunkang10’, *Camellia* DASZ, and *Camellia chekiangoleosa*.

**Supplementary Table 29.** Tissue-specific expression of LSGs across eight different tissues in *Camellia sinensis* var. *sinensis* ’Shuchazao’, *Camellia sinensis* var. *assamica* Yunkang10, *Camellia* DASZ, and *Camellia chekiangoleosa*.

**Supplementary Table 30.** Tissue-specific expression of TFs across eight different tissues in *Camellia sinensis* var. *sinensis* ’Shuchazao’, *Camellia sinensis* var. *assamica* ’Yunkang10’, *Camellia* DASZ, and *Camellia chekiangoleosa*.

**Supplementary Table 31.** Predicted subcellular localization of LSGs and TFs in 13 Theaceae plants.

